# Impact of acetate on CO_2_ fixation pathways in thermophilic and hydrogenotrophic bacteria

**DOI:** 10.1101/2025.03.31.646503

**Authors:** Yoko Chiba, Tomomi Sumida, Masafumi Kameya, Yuto Fukuyama, Tomoyuki Wakashima, Shigeru Shimamura, Ryoma Kamikawa, Yoshito Chikaraishi, Takuro Nunoura

**Affiliations:** Biofunctional Catalyst Research Team, RIKEN Center for Sustainable Resource Science, Japan; Institute for Extra-cutting-edge Science and Technology Avant-garde Research (X-star), Japan Agency for Marine-Earth Science and Technology, Japan; Faculty of Life and Environmental Sciences, University of Tsukuba, Japan; Research Center for Bioscience and Nanoscience (CeBN), Japan Agency for Marine-Earth Science and Technology, Japan; Graduate School of Agricultural and Life Sciences, The University of Tokyo, Japan; Collaborative Research Institute for Innovative Microbiology, The University of Tokyo, Japan; Graduate School of Science and Technology, University Tsukuba, Japan; Graduate School of Agriculture, Kyoto University, Japan; Institute of Low Temperature Science, Hokkaido University, Japan

## Abstract

The Wood-Ljungdahl (WL) pathway and reductive tricarboxylic acid (rTCA) cycle are the dominant chemolithotrophic CO_2_ fixation pathways in bacteria inhabiting aphotic geothermal and deep-sea hydrothermal ecosystems. However, the activity of these bacterial metabolic systems in ecosystems with available organic carbons remains unclear. Here, we examined the impact of extracellular acetate on the CO_2_-fixation pathways of three thermophilic hydrogen-oxidizing and non-acetogenic bacteria using ^13^C tracer-based metabolomics. Under chemolithoautotrophic conditions, *Thermodesulfatator indicus* and *Hydrogenobacter thermophilus* fixed CO_2_ through the WL pathway and rTCA cycle, respectively, whereas *Thermovibrio ammonificans*, which has been suggested to operate both of these pathways, exhibited significant CO_2_ fixation through only the rTCA cycle. Under chemolithomixotrophic conditions with acetate, *H. thermophilus* and *T. ammonificans* assimilated both CO_2_ and acetate via the rTCA cycle. In contrast, acetate suppressed CO_2_ fixation through the WL pathway in *T. indicus* and was used as the primary carbon source under chemolithomixotrophic conditions. These results suggest that the contribution of the WL pathway for CO_2_ fixation might be overestimated in ecosystems where acetate is available. Moreover, the present findings indicate that simultaneous CO_2_ fixation through both the WL pathway and rTCA cycle in a cell, which has been proposed as a possible metabolic strategy for CO_2_-fixation in ancestral life, is not advantageous in extant microorganisms.

## Introduction

Chemolithoautotrophic primary production is essential to sustain aphotic geothermal and deep-sea hydrothermal ecosystems [1], which are recognized as modern analogues of early ecosystems on Earth [2–4]. The Wood-Ljungdahl (WL) pathway and reductive tricarboxylic acid (rTCA) cycle and, the former of which is also called the reductive acetyl-CoA pathway, are the dominant bacterial CO_2_ fixation pathways in these geothermal and hydrothermal environments [1]. However, as the WL pathway and rTCA cycle are fundamentally reversible, it remains unclear if these pathways are capable of CO_2_ fixation in ecosystems where extracellular organic carbons are readily available [5].

Most extant bacterial WL pathways are found in acetogens that utilize CO_2_ as both an electron acceptor and carbon source [6,7], and direct observation of this pathway in non-acetogens is limited. One exception is *Desulfotomaculum acetoxidans*, a mesophilic bacterium isolated from piggery waste that has a functional WL pathway which operates both reductively and oxidatively using hydrogen and acetate, respectively, as electron donors [8–10]. Putative genes for WL pathway enzymes have also been identified in hydrogenotrophic sulfate reducers, including members of the class *Thermodesulfobacteria*, [11,12] suggesting that thermophilic, chemolithoautotrophic non-acetogens play a significant role as primary producers through operation of the WL pathway for CO_2_ fixation in geothermal and hydrothermal environments [11,12]. However, WL pathway function has not been demonstrated in obligately hydrogenotrophic and thermophilic non-acetogens under chemolithoautotrophic or chemolithomixotrophic conditions.

The rTCA cycle is a variant of the TCA cycle, which is also called the citric acid cycle or Krebs cycle [13]. Conventionally, the direction of TCA cycle reactions is predicted by key enzymes in this pathway, particularly those catalyzing the cleavage of citrate (Table S1) [14,15]. Citrate cleavage in the rTCA cycle is generally thought to be catalyzed by ATP-citrate lyase (ACL) [16,17] or by citryl-CoA synthestase (CCS) and citryl-CoA lyase (CCL) in a two-step reaction [18,19]. Although these enzymes have been recognized as signatures of the rTCA cycle, multiple lines of evidence has revealed that the TCA cycle direction is controlled not only by the key enzyme(s), but also by various external and internal factors, including available carbon sources, partial pressure of CO_2_, and amount and/or kinetic parameters of enzymes involved in the cycle [15,20–23]. For instance, *Thermosulfidibacter takaii* [22,24], *Desulfurella acetivorans* [15,23] and other thermophilic and hydrogen-oxidizing bacteria (23) isolated from hydrothermal or geothermal systems operate the TCA cycle in the reductive direction using ATP-independent citrate synthase (CS) at chemolithoautotrophic growth conditions with high-partial pressures of CO_2_. Notably, the TCA cycle operates partially or completely oxidatively when *T. takaii* is grown chemolithomixotrophically with organic acids as carbon sources under high-CO_2_ partial pressure [22]. As another example, the mesophilic sulfate reducer *Desulfobacter postgatei* operates the TCA cycle oxidatively with ACL to generate ATP via the oxidation of acetate [17].

The WL pathway and rTCA cycle are considered to be ancestral forms of CO_2_ fixation systems among the seven known CO_2_ fixation pathways [25–31]. In addition, the occurrence of both the WL pathway and rTCA cycle in a cell is proposed as a possible ancestral form of CO_2_ fixation system based on comparative genomic analyses [28,32,33]. In fact, chemolithoautotrophs that fix CO_2_ through the WL pathway likely also operate at least part of the TCA cycle to obtain the essential cellular metabolic building blocks [28] other than acetyl-CoA, including pyruvate, oxaloacetate (OAA), succinyl-CoA (or succinate), and 2-oxoglutarate (2-OG). However, cooperation of the WL pathway and complete TCA cycle in either working direction has not been demonstrated in any cellular system to date. Although combined genomic and proteomic analyses in the thermophilic hydrogen-oxidizer *Thermovibrio ammonificans* [34] identified components for both the WL pathway and rTCA cycle, a gene encoding a key enzyme in the WL pathway, acetyl-CoA synthetase, was absent [32].

Furthermore, *in-silico* kinetic simulations suggest that increased acetyl-CoA influx caused by a functioning WL pathway would negatively impact the reductive operation of the TCA cycle with CS [35]; however, the ability of the WL pathway and rTCA cycle to operate with ACL has not been theoretically examined.

Members of class *Thermodesulfobacteria* in the phylum *Thermodesulfobacteriota* and classes *Aquificia* and *Desulfurobacteriia* in the phylum *Aquificota* are hydrogenotrophic, extremely thermophilic (optimal growth temperature exceeds 70 □), and comprise the dominant bacterial populations in geothermal and hydrothermal ecosystems. *Thermodesulfobacteria* bacteria are predicted to harbor the WL pathway based on gene homology [11], and *Aquificota* bacteria have a functional rTCA cycle for inorganic carbon fixation [36]. Moreover, *T. ammonificans* in *Desulfurobacteriia* is thought to operate both the WL pathway and rTCA cycle, as described above [32]. Here, we examined the activity of these CO_2_ fixation pathways in *Thermodesulfatator indicus* in *Thermodesulfobacteriota* [12], and *T. ammonificans* and *Hydrogenobacter thermophilus* in *Aquificia* [37–39], under chemolithoautotrophic and chemilithomixotrophic conditions using ^13^C tracer-based metabolomics [22] to evaluate their operation in geothermal and hydrothermal ecosystems and to gain insight into the origin and evolution of central carbon metabolism.

## Materials and Methods

### Strains and growth conditions

*T. indicus* JCM 11887^T^ (=CIR29812), *T. ammonificans* JCM 12110 ^T^ (=HB-1), and *H. thermophilus* JCM 7687 ^T^ (=TK-6) were provided by RIKEN BRC through the National BioResource Project of the MEXT, Japan. Glass test tubes with volumes of 17 and 20 mL (φ16×125 and φ16×150 mm, IWAKI, Japan) were used for the cultivation of *T. ammonificans* and *T. indicus*, respectively, and a 100-mL glass serum bottle (φ40.5×128.0 mm, Maruemu, Japan) was used to cultivate *H. thermophilus*.

*T. indicus*, *T. ammonificans* and *H. thermophilus* were cultivated in inorganic medium (3, 4, and 5 mL, respectively) with H_2_ and CO_2_th as energy and carbon sources, respectively at 70 °C. See the Supporting Information for details.

### Isotopologue and isotopomer analyses

All ^13^C-labeled regents (purity of >99%) used in this study were purchased from Cambridge Isotope Laboratories (Tewksbury, MA, USA).

See the Supporting Information for details on the cell cultivation with ^13^C compounds. To examine CO_2_ incorporation in *T. indicus* cells grown chemolithoautotrophically, cells were harvested 7 h after the addition of 10% (v/v of gas phase) ^13^CO_2_ during the late exponential phase. To examine the incorporation of CO_2_ in *T. indicus* cells grown chemolithomixotrophically with acetate, ^12^C sodium acetate (2 mM final concentration) and 2 mL of ^13^CO_2_, which corresponded to 5% v/v of the final gas phase (H_2_:CO_2_) and 25% of all CO_2_ in the test tube, was added prior to the cultivation. To examine the incorporation of acetate and formate in *T. indicus* cells, [1,2-^13^C_2_] sodium acetate or ^13^C sodium formate (2 mM final concentration), respectively, was added to the inorganic medium prior to cultivation. To examine CO incorporation in *T. indicus* cells, 1 mL of ^13^CO was added to the gas phase to give a final CO:CO_2_ ratio of 1:3.

To examine CO_2_ incorporation in *T. ammonificans* cells grown chemolithoautotrophically, cells in the late exponentially phase were harvested 3 h after the addition of 10% ^13^CO_2_ (v/v of the final gas phase). To examine ^13^CO incorporation in *T. ammonificans* cells, 2.4 mL of ^13^CO was added to the H_2_ atmosphere and a gas mixture of 80% H_2_ and 20% CO_2_ was added to 0.20 MPa. After a 16-h incubation, the cells were harvested by centrifugation. To examine acetate incorporation in *T. ammonificans* cells grown chemolithomixotrophically, [1,2-^13^C_2_] sodium acetate (10 mM final concentration) was added to the inorganic medium prior to cultivation.

To obtain labelled amino acids from *H. thermophilus* cells grown chemolithoautotrophically or chemolithomixotrophically, 10 mL of ^13^CO_2_, which corresponded to ∼10% (v/v) of the gas phase, was added to exponential phase cultures approximately 6 h after inoculation. Cells were harvested by centrifugation 2.5 h after the addition of ^13^CO_2_. To examine the incorporation of acetate in *H. thermophilus* cells, [1,2-^13^C_2_] sodium acetate (10 mM final concentration) was added to the inorganic medium prior to cultivation, and cells were harvested by centrifugation in the late exponential phase.

The harvested cells were stored at −80 °C until needed for further processing. To prepare protein-derived amino acids, the cells were hydrolyzed in 12N HCl at 110 °C and examined using a ZipChip capillary electrophoresis (CE) system (908 Devices, Boston, MA, USA) coupled with an Orbitrap Fusion Tribrid mass spectrometer (Thermo Fisher Scientific, Waltham, MA) as described previously [40]. The obtained MS data was analyzed using Qual Browser in Xcalibur (version 4.3.73.11). The relative abundance of isotopologues and isotopomers of each amino acid was determined from CE-MS- and CE-MS/MS-based data, respectively.

## Results

### Predicted metabolic pathways of the strains used in this study

The distribution patterns of genes encoding WL pathway and TCA cycle enzymes varied amongst the genomes of the three tested species. *T. indicus* encoded a complete set of WL pathway genes, whereas the TCA cycle gene set appeared to lack a gene for the enzymatic interconversion between 2-OG and OAA (Fig. 1A) (see also the Supplementary Text for details, Fig. S1 and Tables S2 and S3). *T. ammonificans* possessed all of the genes for the rTCA cycle, and also harbored most of the candidate genes for WL pathway enzymes, with the exception of acetyl-CoA synthetase (Figs. 1A and S1) [32]. *H. thermophilus*, which serves as a model organism for understanding the machinery of the rTCA cycle [41–45], lacked most of the genes related to the WL pathway (Figs. 1A and S1) [32].

**Fig. 1.**
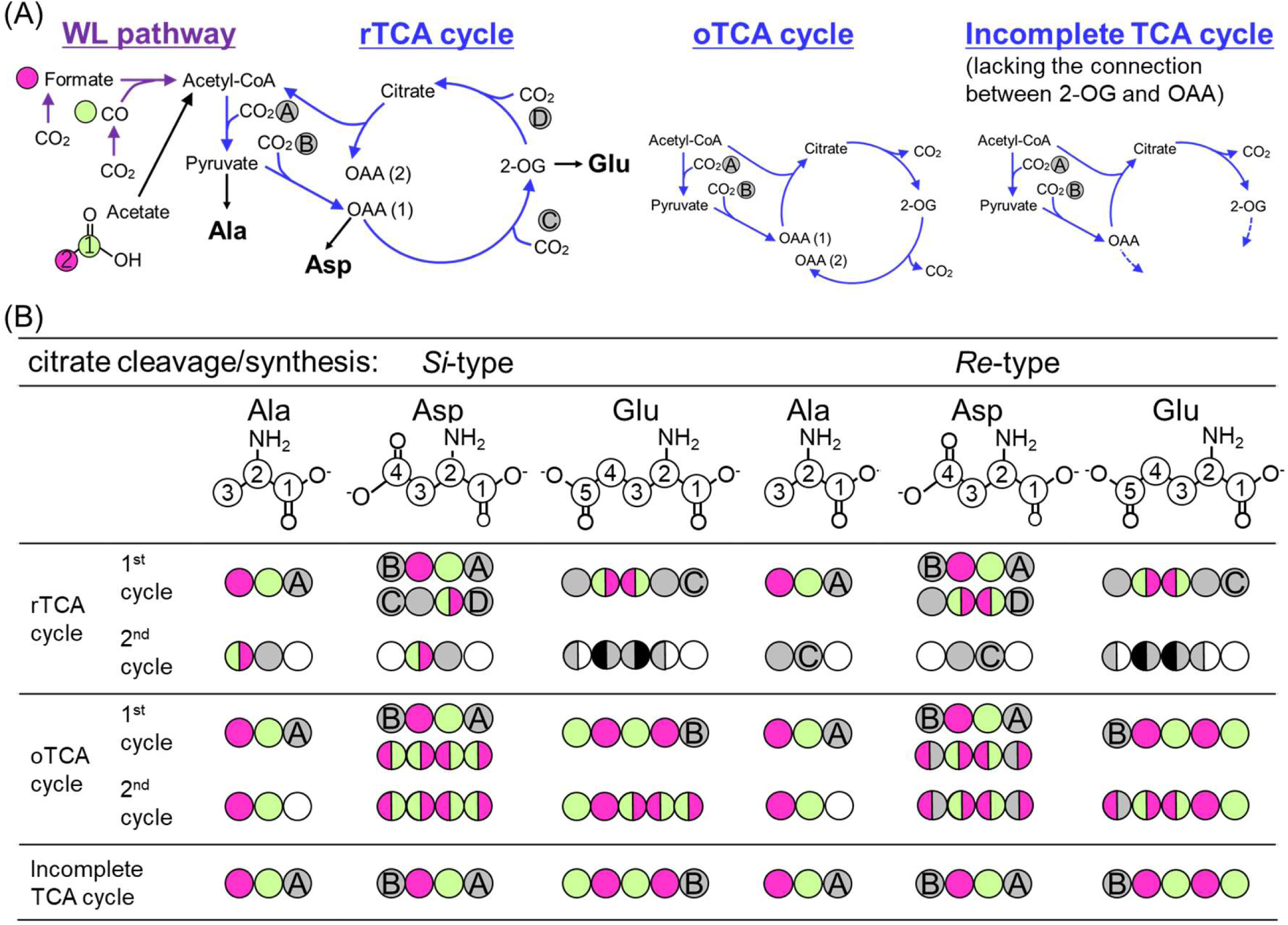
Schematics of the WL pathway and reductive, oxidative and incomplete TCA cycles (A), and the expected labeling patterns of metabolites (B). (A) Reactions included in the WL pathway and TCA cycle are shown with purple and blue arrows, respectively. rTCA cycle: reductive TCA cycle; oTCA cycle: oxidative TCA cycle; OAA: oxaloacetate; 2-OG: 2-oxoglutarate. (B) Carbons derived from CO, formate, and acetate are shown with light green, magenta and light green (C1 of acetate) and magenta (C2 of acetate), respectively. Carbons fixed through the first cycle of the TCA cycle are highlighted in gray and labeled from “A” to “D” based on the reactions shown in (A). Black circles indicate carbons derived from CO or formate. Note that the labeling pattern differs based on the stereochemistry of the citrate cleavage/synthesis enzymes. Upper and lower Asps are derived from OAA (1) and OAA (2), respectively, in (A).

Among the three examined strains, variations in the gene involved in citrate synthesis/cleavage reactions were also identified. CS is classified into *Si*- and *Re*-types based on the stereospecificity of the binding manner of acetyl-CoA and OAA (Fig. S2), and the reactions catalyzed by these enzymes can be determined by isotopomer structure using a tracer-based metabolomics approach (Fig. 1B) [46,47]. *T. indicus* has a homolog of *Re*-CS (Fig. S3; see also details in the Supplementary Text), whereas *T. ammonificans* and *H. thermophilus* harbor ACL and CCS/CCL, respectively, and lack CS (Table S2). CCL is the ancestral form of ACL and *Si*-CS [48,49] and predicted to have the same stereochemistry as *Si*-CS [23,45]. All the three strains have candidate gene for acetyl-CoA synthase, which catalyzes the conversion of acetate to acetyl-CoA (Fig. S1).

### Effects of acetate on growth and protein expression by the three hydrogen oxidizers

*T. indicus, T. ammonificans*, and *H. thermophilus* were cultivated chemolithoautotrophically with H_2_ and CO_2_ as energy and carbon sources, respectively, and also chemolithomixotrophically using acetate as a carbon source in addition to CO_2_ under the hydrogenotrophic condition. Under chemolithomixotrophic conditions, the growth of *T. indicus* and *H. thermophilus* was slightly stimulated and the final cell density was increased by 20% compared to that obtained under chemolithoautotrophic conditions (Fig. S4), as reported previously [12,50]. In contrast, the addition of acetate had no discernible effect on the growth rate or final cell density of *T. ammonificans* (Fig. S4).

Gene expression related to the WL pathway and TCA cycle was also examined by shotgun proteomics. In *T. indicus*, all of the enzymes in the WL pathway and an incomplete set of TCA cycle enzymes were expressed under chemolithoautotrophic and chemolithomixotrophic conditions (Table S4). In *H. thermophilus*, all TCA cycle enzymes, except for a few subunits of heteromeric proteins, were detected under both chemolithoautotrophic and chemolithomixotrophic conditions (Table S5). The expression of both WL pathway and TCA cycle proteins in *T. ammonificans* under chemolithoautotrophic condition has been reported previously [32].

### Carbon flow in *T. indicus*

The operation and working direction of the WL pathway and/or TCA cycle were examined in all three species using tracer-based metabolomics. Tables 1 and 2 summarize the pattern of isotopomers and isotopologues of the amino acids, Ala, Asp and Glu, which were obtained by protein hydrolysis in cells grown with ^13^C-labeled substrates. The patterns of Ala, Asp and Glu reflected those of their precursors: pyruvate, OAA and 2-OG, respectively [40].

*T. indicus* cells grown chemolithoautotrophically with ^13^CO_2_ incorporated single or multiple ^13^C molecules in both the C-1 and non-C-1 positions of Ala and Asp (Table 2). In contrast, when ^13^CO or [^13^C] formate was added together with ^12^CO_2_ in the culture medium, incorporation of single ^13^C molecules into non-C-1 positions was observed in most labeled Ala and Asp (Table 2). These results were consistent with the expected labeling patterns, which are based on the assumption that CO_2_, CO, and formate are directly incorporated into acetyl-CoA through the WL pathway, and that acetyl-CoA is carboxylated to pyruvate and then further carboxylated to form OAA (Fig. 1A, Fig. S1). The oxidation of CO to CO_2_ was negligible, whereas ∼ 5% of the formate incorporated into the cells was oxidized to CO_2_ (complete experimental details and results are provided in the Supplementary Text and Fig. S5). Taken together, these results demonstrate that the WL pathway functions as a chemolithoautotrophic CO_2_ fixation pathway in the extremely thermophilic and non-acetogenic bacterium *T. indicus*.

The working direction of the TCA cycle in *T. indicus* was next assessed based on the labeling pattern of Glu and the following rationale. If 2-OG is synthesized from OAA through the rTCA cycle, no ^13^C would be observed in the C-1 position of Glu when *T. indicus* is cultivated with ^13^CO or [^13^C] formate in cases when the CO and formate are not oxidized to CO_2_ (Fig. 1B, rTCA cycle with *Re*-type citrate cleavage/synthesis; Fig. S6). In contrast, if 2-OG is synthesized through the oxidative TCA (oTCA) cycle, the C-1 position of Glu would be labeled with ^13^C derived from ^13^CO, but not from [^13^C] formate (Fig. 1B, Fig. S7). We found that the ratio of ^13^C_1_ Glu with ^13^C in the C-1 position was 5.1% and 2.1% in *T. indicus* cells cultured in the presence of ^13^CO and [^13^C] formate, respectively (Table 2). The low percentage of ^13^C_1_ Glu with ^13^C in the C-1 position in cultures with [^13^C] formate was likely due to the incorporation of CO_2_ derived from oxidized [^13^C] formate. These findings suggest that 2-OG was synthesized through the oTCA cycle. Notably, however, the connection and working direction among OAA, succinyl-CoA and 2-OG could not be determined based on the isotopomer patterns obtained in this study.

The impact of acetate on the WL pathway was also analyzed under chemolithomixotrophic conditions. If all cellular acetyl-CoA is synthesized from CO_2_ through the WL pathway, it is expected that in the presence of ^13^CO_2_ and ^12^CO_2_, the ratio of ^13^C_1_ Ala, which is derived from pyruvate through acetyl-CoA, with ^13^C at the C-1 position to the C-2 or C-3 position would be 1:2. In *T. indicus* cells grown chemolithoautotrophically under these conditions, the observed ratio was nearly 1:2 (8.4:17.6). In contrast, in cultures supplemented with non-labeled acetate and ^13^CO_2_ (^13^CO_2_: ^12^CO_2_ =1:3), the ratio was altered to approximately 4:1 (24.5:6.2; Table 2), indicating that more than 85% of acetyl-CoA was derived from the external acetate source. Considering that the cell yield under chemolithomixotrophic conditions with acetate increased by only 20% compared to the yield resulting from chemolithoautotrophic growth, CO_2_ fixation through the WL pathway was likely suppressed by the increased acetyl-CoA influx from acetate.

The working direction of the TCA cycle in *T. indicus* cells grown chemolithomixotrophically with [1,2-^13^C_2_] acetate and ^12^CO_2_ was next investigated. Under these conditions, 37.3% of Glu was in the form of ^13^C_4_, with the C-1 carboxyl group being labeled with ^13^C (Table 2). The observed labeling pattern confirmed that 2-OG was synthesized with acetyl-CoA and OAA through oxidative operation of the TCA cycle (Fig. 1B, see the Supporting Information for details). The labelling patterns of Asp (49.9% ^13^C_2_ and 0.3% ^13^C_3_) indicated that the majority of OAA was synthesized through the direct carboxylation of pyruvate, but not from ^13^C_4_ 2-OG via the oTCA cycle. Collectively, these results reveal that the TCA cycle is incomplete and bifurcated in *T. indicus*, and that enzymatic reactions between 2-OG and OAA do not occur under autotrophic or mixotrophic conditions, as would be expected based on the genome sequence (Fig. 1 A and Fig. S8).

### Carbon flow in *T. ammonificans*

When *T. ammonificans* was cultivated with ^13^CO and non-labeled CO_2_, no significant incorporation of ^13^C in Ala, Asp and Glu was observed, in contrast to the case of *T. indicus* (Fig. 2, Table 1), indicating that the WL pathway had a negligible contribution to CO_2_ fixation. Under chemolithoautotrophic conditions, the labeling patterns of *T. ammonificans* cells grown with ^13^CO_2_ and ^12^CO_2_ (1:2) were consistent with those expected by operation of the rTCA cycle with ACL. Namely, the observed abundance of ^13^C_2_ and ^13^C_3_ Glu (17.7% and 4.9%, respectively) (Table 2) suggests that 2-OG was only synthesized through reductive operation of the TCA cycle, assuming that the proportion of acetyl-CoA derived from the WL pathway was negligible (Fig. 1, Fig. S6A and Fig. S7A). The formation of ^13^C_2_ Ala, which had ^13^C in both C-1 and C-2 or C-3 positions (5.5%), and ^13^C_2_ Asp, which had two ^13^C in positions other than the C-1 carboxyl group (6.3%), can be explained by a second turn of the rTCA cycle. Moreover, a lower abundance of ^13^C_2_ Ala with ^13^C in both the C-2 and C-3 positions was detected (3.0%), suggesting that this compound may be a signature of the third turn of the rTCA cycle.

**Fig. 2.**
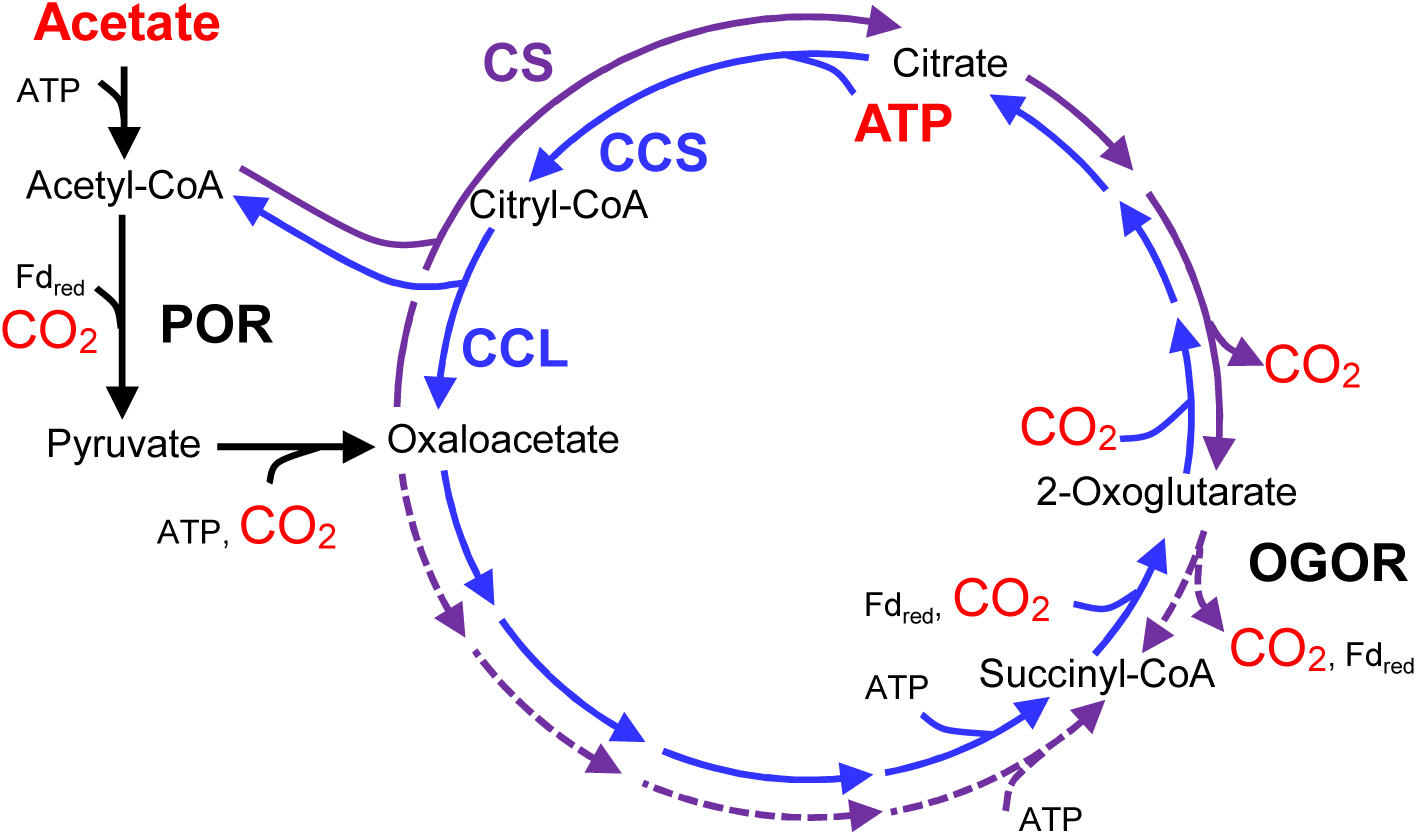
Flow of the TCA cycle under chemolithomixotrophic conditions with acetate in *H. thermophilus* and *T. takaii*. *H. thermophilus* (blue arrows) and *T. takaii* (purple arrows) have ATP-dependent (CCS/CCL) and ATP-independent (CS) citrate cleavage/synthesis systems, respectively. Black arrows indicate common reactions in the two organisms. Dotted arrows indicate the expected working direction of the TCA cycle, which cannot be theoretically determined based on isotopomer patterns. CCS: Citryl-CoA synthetase; CCL: Citryl-CoA lyase; CS: Citrate synthase; POR: pyruvate:ferredoxin oxidoreductase; OGOR: 2-OG:ferredoxin oxidoreductase; Fd_red_: reduced ferredoxin

We also examined the assimilation of [1,2-^13^C_2_] acetate by *T. ammonificans* in the presence of H_2_ and ^12^CO_2_. ^13^C_2_ Ala comprised 10.4% of total Ala, indicating that acetate was incorporated into the cells (Table 1). Approximately 3% of ^13^C excess was observed in ^13^C_1_ Asp and Glu, indicating that the rTCA cycle was functional (Table 1). As acetate had no impact on the growth of *T. ammonificans*, which also exhibited lower acetate assimilation compared to *T. indicus* and *H. thermophilus*, we concluded that *T. ammonificans* passively assimilated acetate and that the rTCA cycle was not significantly affected by the acetyl-CoA influx from the external acetate source. The passive assimilation of acetate by in *T. ammonificans* is consistent with the lack of a candidate gene for a cation/acetate symporter (KEGG ORTHOLOGY: K14393), which was identified in *T. indicus* and *H. thermophilus* (Thein_1042 and HTH_1561, respectively).

### Carbon flow in *H. thermophilus*

The labeling patterns of *H. thermophilus* cells grown with ^13^CO_2_ and ^12^CO_2_ were consistent with those produced by a functional rTCA cycle that includes CCS and CCL, which has the same stereochemistry with that of *Si*-CS (Fig. S6A). The detection of ^13^C_3_ and ^13^C_4_ Glu and ^13^C_2_ Asp with two ^13^C in carboxyl groups located at positions other than C-1 (6.6%) confirmed operation of the rTCA cycle, but not the oTCA cycle (Fig. 1 and Table 1). Further, the production of ^13^C_2_ Ala with ^13^C in both the C-1 and C-2 or C-3 positions (6.7%) would be expected in the second turn of the rTCA cycle (Table 2). The ^13^C labeling patterns with ^13^CO_2_ confirmed that *H. thermophilus* operates the rTCA cycle under the tested chemolithoautotrophic conditions.

The working direction of the TCA cycle in *H. thermophilus* was also examined under chemolithomixotrophic conditions with supplemental acetate. In cells grown with [1,2-^13^C_2_] acetate and ^12^CO_2_, more than 50% of Ala, Asp and Glu contained one or two ^13^C atoms, confirming that *H. thermophilus* actively incorporated acetate into cellular compounds. The production of ^13^C_1_ Ala, Asp and Glu would occur during the second cycle of the rTCA cycle, but would not be formed by the oTCA cycle or an incomplete TCA cycle (Fig. 1). Notably, the amount of ^13^C_4_ Glu, which can be produced only through the oTCA cycle, was negligible despite the high ratio of [1,2-^13^C_2_] acetate incorporation by *H. thermophilus*. The operation of the rTCA cycle was also supported by isotopologue analysis of cells grown with non-labeled acetate and ^13^CO_2_. Under these conditions, the detection of Ala with ^13^C in C-1, Asp with ^13^C in non-C-1, and Glu with ^13^C in C-1 positions indicated the carboxylation of acetyl-CoA, pyruvate, and succinyl-CoA, respectively.

## Discussion

### Impact of acetate on TCA cycle variants

Under chemolithomixotrophic conditions, in which H_2_, acetate and high partial pressures (20%) of CO_2_ are available, the impact of acetate on the TCA cycle, particularly on citrate cleavage/synthesis reactions, varies depending on the types of enzymes catalyzing these reactions. The findings from the present study revealed that the hydrogen-oxidizing bacteria *H. thermophilus* (Fig. 2) and *T. ammonificans* operate the rTCA cycle, including the associated citrate cleavage reaction(s), under chemolithomixotrophic growth conditions in the presence of acetate, as was reported for the green sulfur bacterium *Chlorobaculum tepidum* [51]. Thus, the reductive operation of the TCA cycle with ATP-dependent citrate cleavage reactions in cells that utilize acetate and CO_2_ as carbon sources appears to be a shared strategy among chemolithotrophs and photolithotrophs.

In contrast to the above hydrogen-oxidizing bacteria, citrate synthesis with CS occurs in the obligate hydrogenotroph *T. takaii* (Fig. 2) and facultative hydrogenotroph *D. acetivorans* under chemolithomixotrophic conditions in the presence acetate, whereas their TCA cycle proceeds in the reductive direction with citrate cleavage with CS under chemolithoautotrophic conditions [15,22]. The marked difference in the operation of the TCA cycle under chemolithomixotrophic conditions with acetate can be explained by the thermodynamic properties of the enzymes used for citrate cleavage/synthesis. ATP-dependent citrate cleavage by ACL (*T. ammonificans* and *C. tepidum*) or CCS/CCL (*H. thermophilus*) is thermodynamically close to neutral (ΔG^0’^=5.6 kJ/mol), whereas ATP-independent citrate cleavage with CS (*T. takaii* and *D. acetivorans*) is highly endergonic (ΔG^0’^=37.4 kJ/mol) [17] (Table S1). Thus, citrate cleavage is more likely to occur in *H. thermophilus, T. ammonificans* and *C. tepidum* than in *T. takaii* and *D. acetivorans* due to these thermodynamic differences. The observed differences in the operational direction of the TCA cycle in these species in response to high acetyl-CoA influx due to acetate assimilation can, at least in part, be explained by the thermodynamics of the enzymes involved in citrate cleavage/synthesis.

The activity of ACL with respect to citrate cleavage and synthesis is also affected by the electron donor source. Notably, when the acetate-oxidizing chemoorganoheterotroph *D. postgatei* is cultured with acetate as a carbon source and the sole electron donor, the oTCA cycle functions with ACL, even under a high (20%) CO_2_ partial pressure [52,53]. To operate the rTCA cycle, reduced ferredoxin (Fd_red_) is required for the conversion of succinyl-CoA to 2-OG by 2-OG:ferredoxin oxidoreductase (OGOR) [14] and for acetyl-CoA conversion to pyruvate by pyruvate:ferredoxin oxidoreductase (POR) (Fig. 2). The reduction of acetyl-CoA by POR is also an essential reaction in gluconeogenesis. Although the machinery involved in Fd reduction in the rTCA cycle has not been experimentally proven [15], *H. thermophilus, T. ammonificans* and *C. tepidum* use hydrogen or sulfur compounds as an electron donor to generate Fd_red_. In contrast, *D. postgatei* obtains Fd_red_ by oxidizing acetate when provided as the sole electron donor. The finding that *D. postgatei* oxidizes 2-OG to succinyl-CoA with OGOR [52] to provide Fd_red_, which may be used for the reduction of acetyl-CoA to provide pyruvate for gluconeogenesis, suggests that acetyl-CoA reduction via POR is primarily regulated by the Fd_red_ supply. Because the flux of acetyl-CoA reduction should be smaller than that of 2-OG oxidation, ACL catalyzes citrate synthesis when acetate is the sole electron donor.

### Possible simultaneous operation of the WL pathway and rTCA cycle

If the WL pathway and rTCA cycle are functioning simultaneously for CO_2_ fixation in a cell, acetyl-CoA influx would be increased compared to cells with only one functional pathway [35]. The presence of extracellular acetate also affects acetyl-CoA influx when the acetate is actively incorporated into cells, as this process serves as an analogue of acetyl-CoA influx through CO_2_ fixation by either of the pathways. Therefore, the simultaneous operation of the WL pathway and rTCA cycle can be assessed based on comparisons to cells grown chemolithomixotrophically with acetate and chemolithoautotrophically.

In the present study, we could not find any evidence for the simultaneous operation of the WL pathway and rTCA cycle for CO_2_ fixation in the three examined thermophilic and hydrogenotrophic species. Although combined genomic and proteomic analyses previously suggested that WL pathway is used to fix CO_2_ in *T. ammonificans* [32], the present tracer-based metabolomic analyses of *T. ammonificans* cultured under chemolithoautotrophic conditions did not detect labeled metabolites to support the operation of the WL pathway. Furthermore, the observed growth properties together with the tracer-based metabolomic analyses in *T. indicus* grown chemolithomixotrophically with acetate suggest that the contribution of the WL pathway for acetyl-CoA synthesis was suppressed under high acetyl-CoA influx (Table 1, Fig. S4). This response is distinct from that of *H. thermophilus* grown under chemolithomixotrophic conditions, in which the rTCA cycle continues to operate in the reduction direction with ATP-dependent citrate cleavage enzyme(s). Moreover, previous theoretical analysis [35] suggests that increased acetyl-CoA influx of the WL pathway negatively affects the reductive operation of the TCA cycle with CS, and in fact, the TCA cycle operates oxidatively in the presence of acetate in *T. takaii*. Taken together, these findings suggest that acetyl-CoA production through the WL pathway is suppressed if the rTCA cycle with ACL is active in the same cell, whereas the reductive operation of TCA cycle with CS is suppressed by the WL pathway in the same cell.

No extant organism has been found to possess complete gene sets for both the WL pathway and TCA cycle [32]. Recent tracer-based metabolomic analyses of *Methanothermobacter thermautotrophicus*, which grows chemolithoautotrophically using the WL pathway, have revealed the operation of an incomplete rTCA cycle lacking aconitase and isocitrate dehydrogenase for CO_2_ fixation, as was suggested by previous genomic analyses [40,54]. This type of incomplete rTCA cycle was also predicted based on comparative genomic analysis to function in *Moorella thermoacetica*, which also has a functional WL pathway [55]. These observations also suggest that organisms do not gain an advantage by simultaneously operating the WL pathway and complete rTCA cycle for CO_2_ fixation.

### Effects of acetyl-CoA influx in non-acetogenic chemolithotrophs in hydrothermal ecosystems

The present results suggest that ATP-dependent citrate cleavage is an essential reaction for rTCA cycle activity in geothermal and hydrothermal microbial ecosystems where organic compounds, such as acetate, are available [5]. In contrast, the WL pathway is adversely affected by increased acetyl-CoA influx, and thus, the contribution of the WL pathway for CO_2_ fixation might be overestimated in these ecosystems. However, the present study has also demonstrated that the carboxylation of acetyl-CoA to pyruvate and pyruvate to OAA are not affected by the influx of acetyl-CoA in chemolithotrophs with a functional rTCA cycle or WL pathway. These findings are consistent with previous observations in a hydrogenotrophic methanogen and heterotrophic thermophile. The hydrogenotrophic methanogen *M. thermautotrophicus* utilizes an incomplete rTCA cycle to synthesize pyruvate, OAA and 2-OG through CO_2_ fixation in the presence of acetate [40,56]. The hyperthermophilic and heterotrophic bacterium *Thermotoga neapolitana* also synthesizes pyruvate through the carboxylation of acetyl-CoA in the presence of acetate and high CO_2_ partial pressure [57]. These findings suggest that the contribution of CO_2_ fixation for the synthesis of essential building blocks in both chemolithoautotrophs and chemoorganoheterotrophs may be larger than expected in CO_2_-rich hydrothermal and aphotic geothermal microbial ecosystems, even in the presence of organic compounds. Moreover, they also imply that the importance of CO_2_ fixation and the boundary between autotrophs (or mixotrophs) and heterotrophs in aphotic geothermal and deep-sea hydrothermal ecosystems, as well as the early ecosystems on Earth, is more complicated and blurred than previously thought.

## Conclusion

In summary, the presented results together with those from previous reports have revealed that organic acids, particularly acetate, affect the activity of bacterial carbon fixation systems, including the rTCA cycle and WL pathway, which are the dominant CO_2_ fixation systems in aphotic hydrothermal and geothermal microbial ecosystems. Although the impact of increased acetyl-CoA influx in the presence of extracellular acetate is limited in the rTCA cycle, which utilizes ATP-dependent citrate cleavage systems for CO_2_ fixation, such influx affects the operational direction of the reversible TCA (roTCA) cycle, which functions with

ATP-independent citrate cleavage enzymes (CS) (15, 22). CO_2_ fixation through the WL pathway is also markedly reduced by increased acetyl-CoA influx, and thus, the contribution of the WL pathway in non-acetogenic chemolithotrophs for primary production is potentially overestimated in hydrothermal and aphotic geothermal microbial ecosystems with available organic acids [1].

## Acknowledgement

This work was supported in part by JSPS KAKENHI Grant Number 19K15745 and 23H04654 (YC),19H00988 and 19H05684 (TN), the Institute for Fermentation, Osaka (IFO) research grant G-2024-1-011 (RK) and The Green Innovation Fund Project, JPNP22010, commissioned by the New Energy and Industrial Technology Development Organization (NEDO). We appreciate the fruitful discussions of Dr. Donate Giovanelli and Dr. Constantino Vetriani regarding the metabolism of *T. ammonificans* prior to starting the study. We also thank Dr. Sanae Sakai and Nao Tsunematsu for assistance with microbial cultivation, Dr. Eiji Tasumi for help with developing a cultivation system, and Masayuki Miyazaki for preliminary organic acid measurements for tracer-based metabolomics.

## Data Availability Statement

The datasets generated during and/or analyzed during the present study are available from the corresponding author on request.

---

**Fig. S1.**
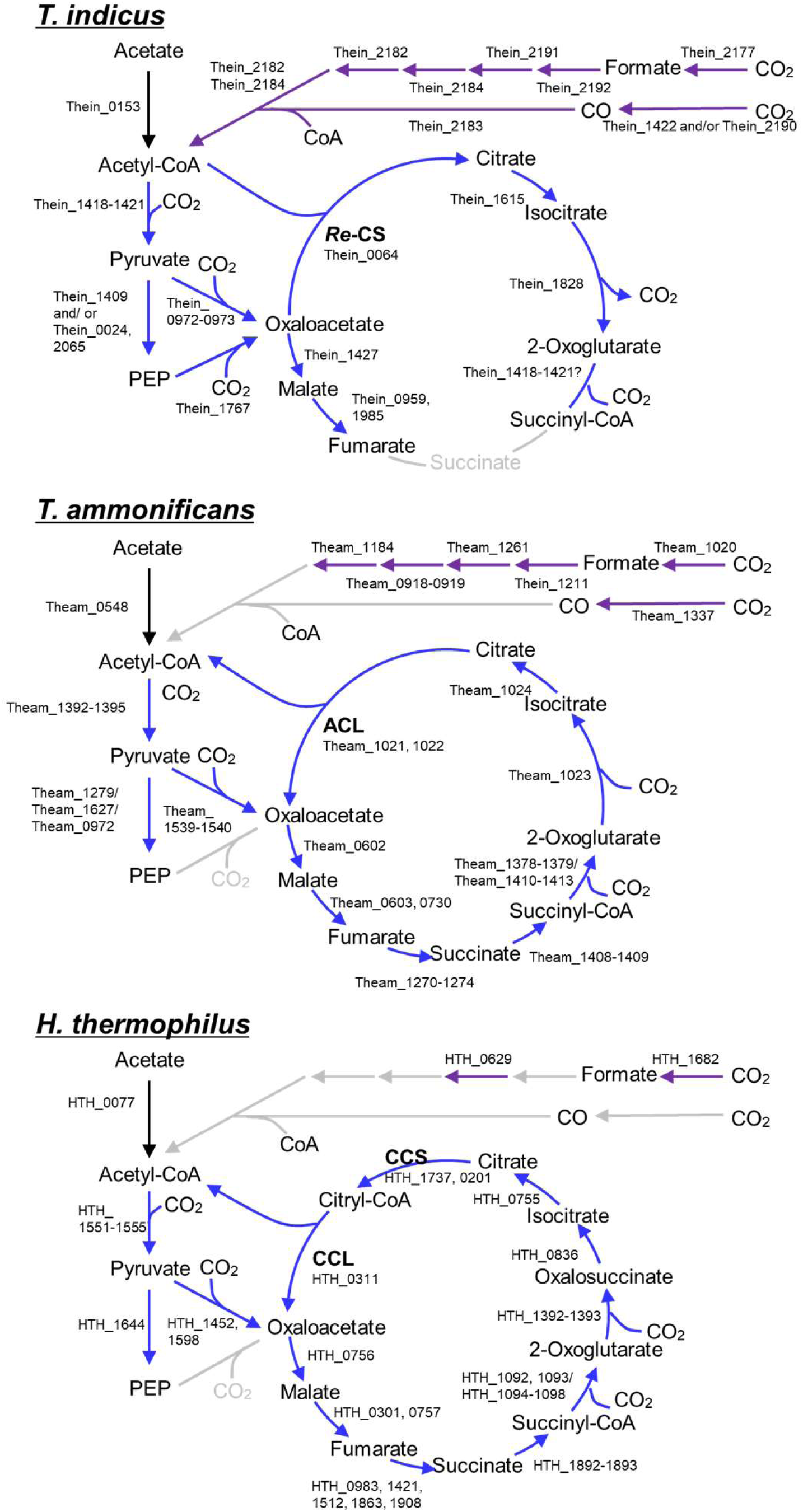
CO_2_ fixation pathways predicted from comparative genomic analysis. Purple and blue arrows indicate reactions related to the WL pathway and TCA cycle, respectively. The gene IDs for the predicted proteins in each pathway are shown beside each arrow. Gray arrows indicate that the candidate gene is absent. The data suggests that *T. indicus* uses the WL pathway, whereas *T. ammonificans* and *H. thermophilus* utilized the rTCA cycle for CO_2_ fixation.

**Fig. S2.**
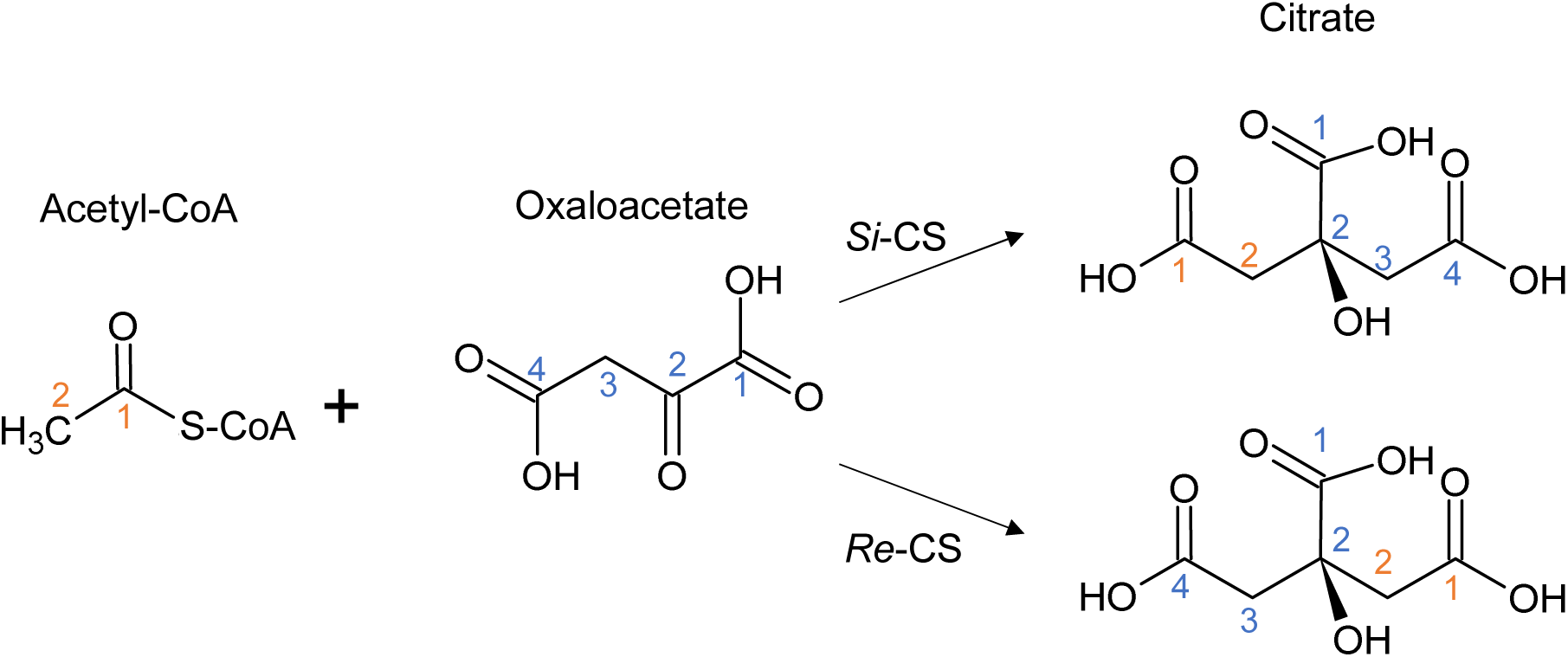
Stereochemistry of *Si*- and *Re*-citrate synthases (CS). ATP-citrate lyase is expected to have the same stereochemistry as *Si*-CS based on high amino acid sequence homology.

**Fig. S3.**
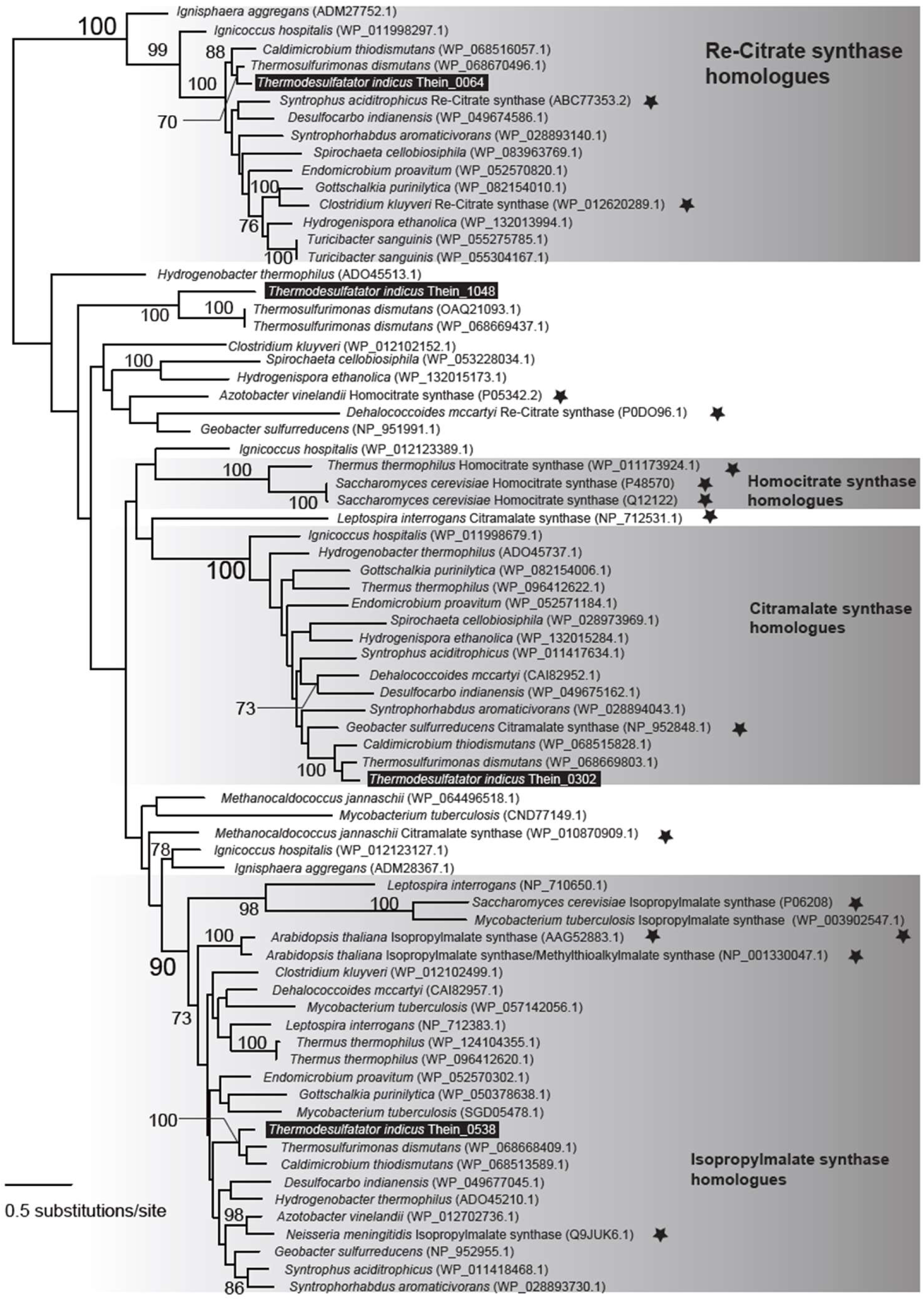
Phylogenic tree of *Re*-CS, isopropylmalate synthase, and their paralogues. Proteins in *T. indicus* are shown with white letters in black boxes. Proteins whose function have been biochemically analyzed are highlighted with stars. This tree was reconstructed by the maximum likelihood framework. Maximum likelihood bootstrap values larger than 70% are shown on branches. Homologues of *Re*-CS, isopropylmalate synthase, citramalate synthase, and homocitrate synthase from prokaryotes and eukaryotes were retrieved from the non-redundant protein database of GenBank using amino acid sequences of *T. indicus* homologues and those of proteins biochemically analyzed (highlighted by stars) as queries. Those protein sequences were aligned with the L-INS-i of MAFFT [62], followed by removal of ambiguously aligned sites using BioEdit [63], resulting in the final dataset, which comprised 74 taxa and 332 sites. The final dataset was subjected to phylogenetic analysis by IQ-Tree v. 1.6.7 [64] with 100 bootstrap analyses under the LG+G+F model.

**Fig. S4.**
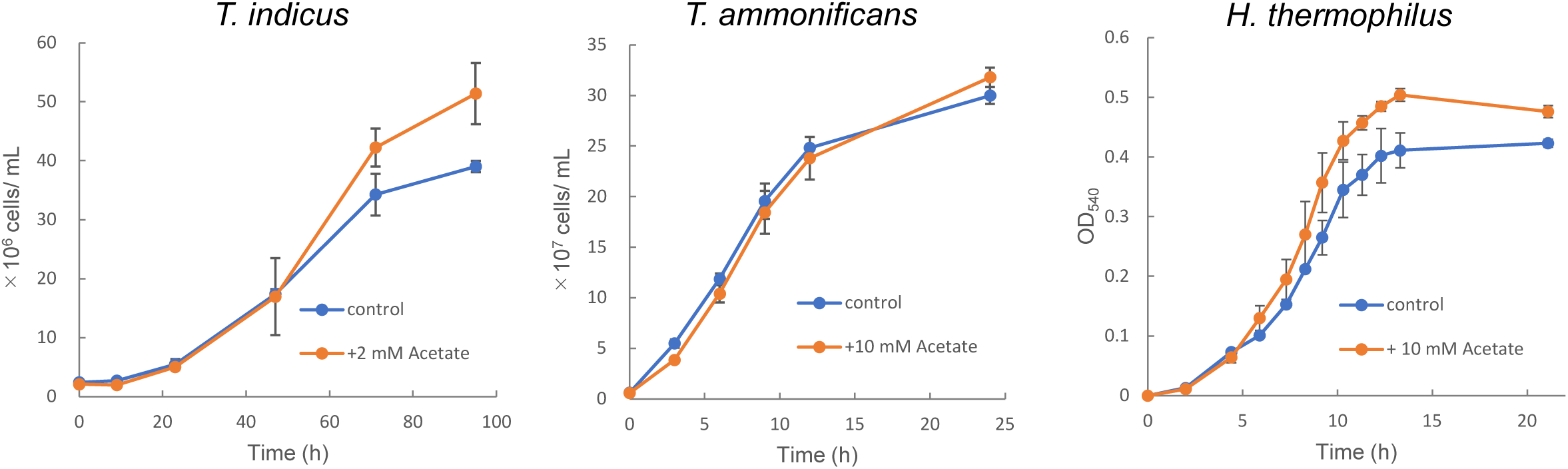
Effect of supplemental acetate on the growth of *T. indicus*, *T. ammonificans* and *H. thermophilus*. The growth of these three species were compared between chemolithoautotrophic (light blue) and chemolithomixotrophic conditions (orange) with the indicated concentration of sodium acetate. Error bars indicate the standard division.

**Fig. S5.**
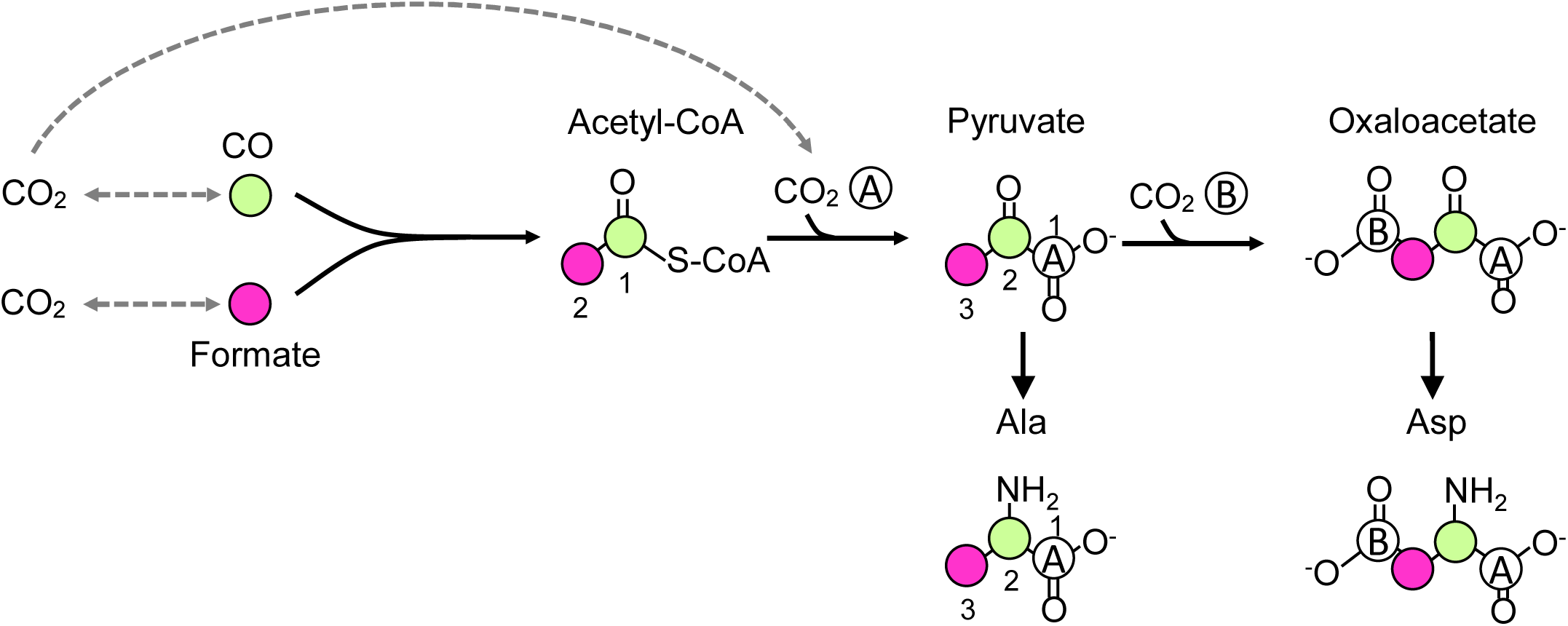
Incorporation of CO and formate via the WL pathway. If CO and formate (light green and magenta circles, respectively) are directly fixed into acetyl-CoA through the WL pathway, they would compose the C-1 and C-2 carbons, respectively, of the acetyl moiety of acetyl-CoA. Acetyl-CoA is expected to be carboxylated to pyruvate, and pyruvate is further carboxylated to oxaloacetate. Therefore, if C-1 of Ala is labeled with ^13^C in cells cultivated with ^13^CO or ^13^C-formate and ^12^CO_2_, it indicates that the ^13^C-labeled CO and/or formate were oxidized to CO_2_ and then incorporated into cells, as shown by the dotted gray arrows.

**Fig. S6.**
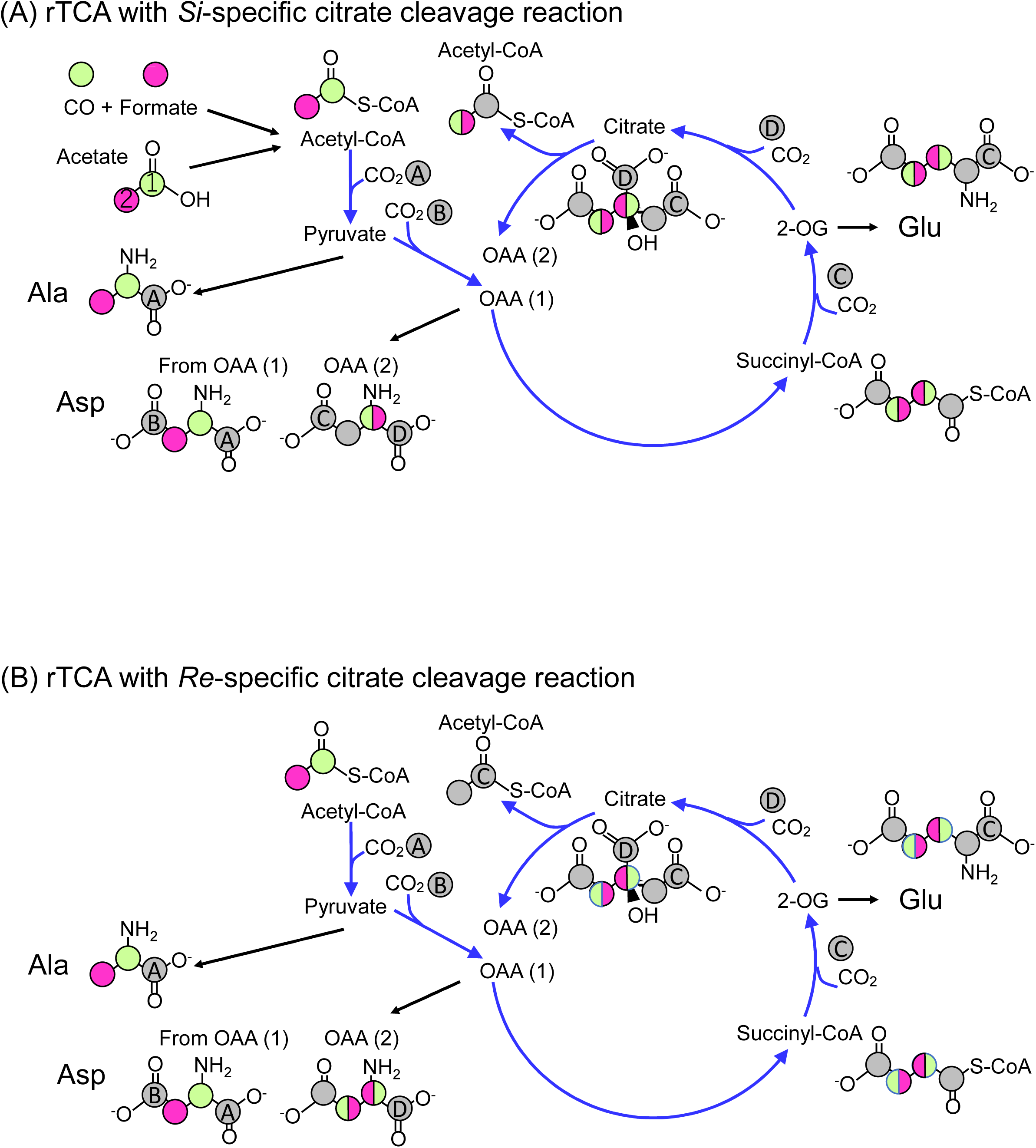
Expected ^13^C-labeleling patterns of metabolites of the rTCA cycle with *Si*-specific (A) or *Re*-specific (B) citrate cleavage reactions. Carbons originating from CO, formate and acetate are shown in light green, magenta, and light green or magenta, respectively. Carbons fixed through the TCA cycle are highlighted in gray and labeled from “A” to “D”.

**Fig. S7.**
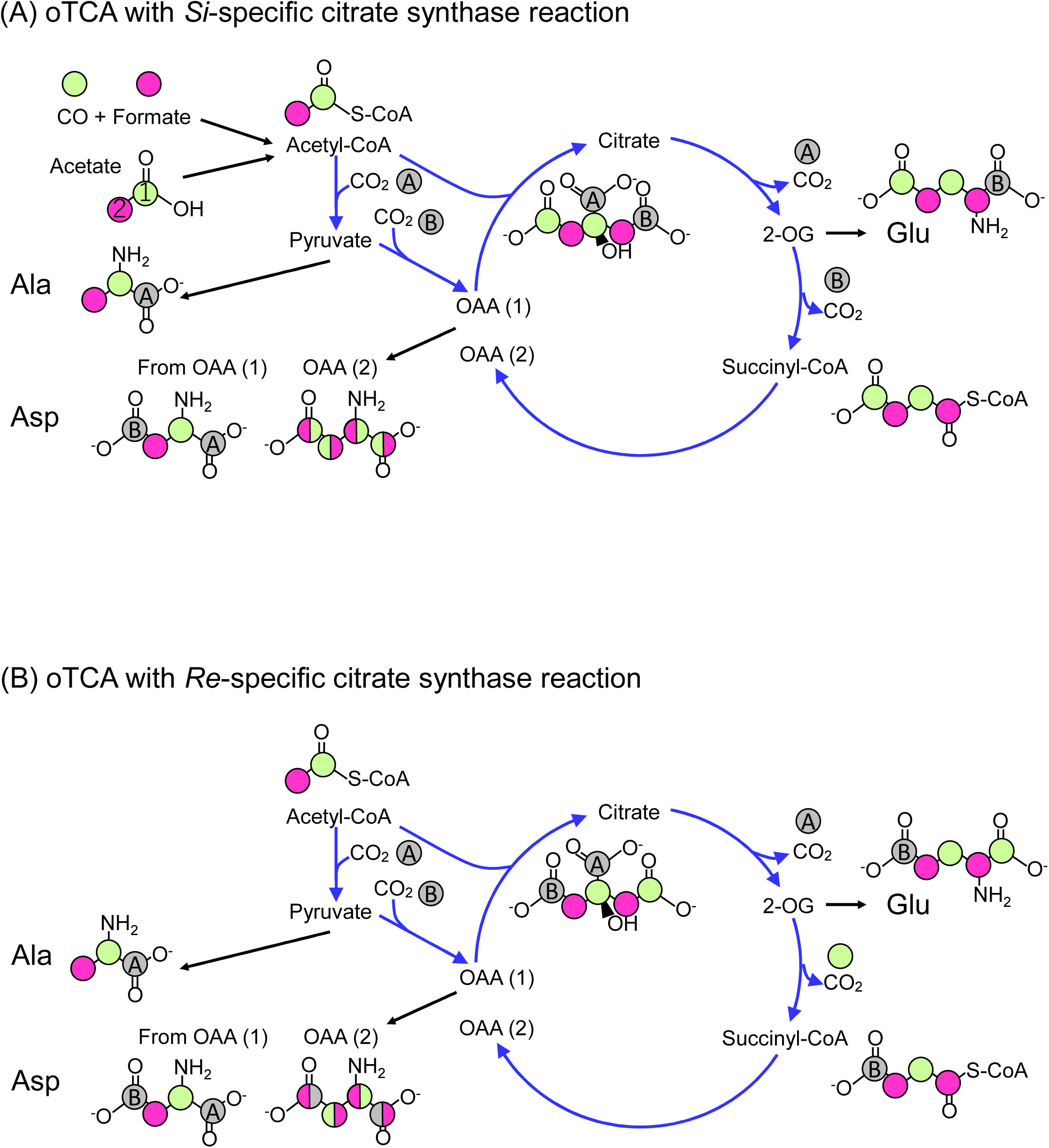
Expected ^13^C-labeleling patterns of metabolites of the oTCA cycle with *Si*-specific (A) or *Re*-specific (B) citrate synthesis reactions. Carbons originating from CO, formate and acetate are shown in light green, magenta, and light green or magenta, respectively. Carbons fixed and released through the TCA cycle are highlighted in gray and labeled “A” and “B”.

**Fig. S8.**
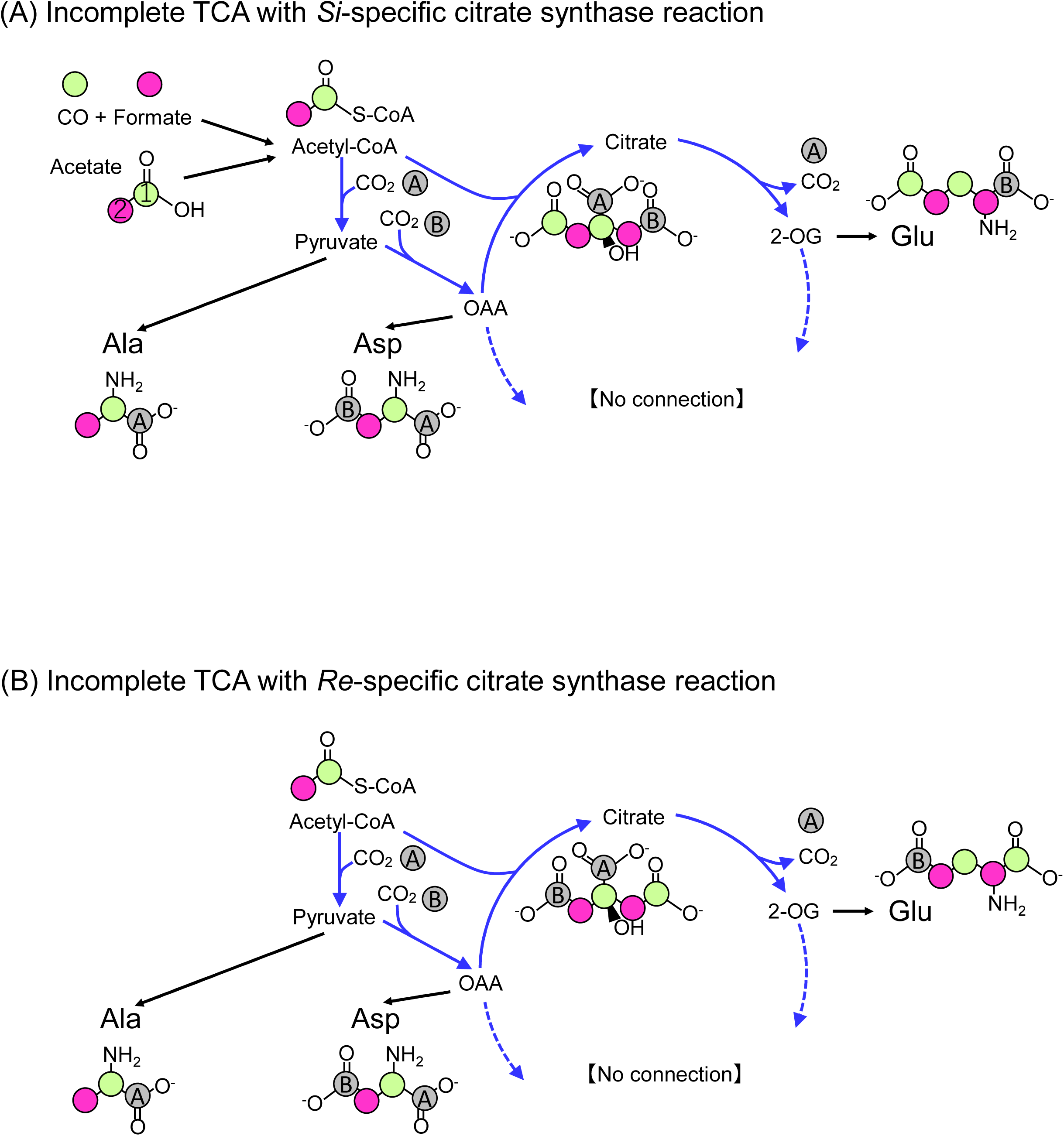
Expected ^13^C-labeleling patterns of metabolites of the incomplete TCA cycle with *Si*- specific (A) or *Re*-specific (B) citrate synthesis reactions. Carbons originating from CO, formate and acetate are shown in light green, magenta, and light green or magenta, respectively. Carbons fixed and released through the TCA cycle are highlighted in gray and labeled “A” and “B”.

## References

1. Dick GJ. The microbiomes of deep-sea hydrothermal vents: distributed globally, shaped locally. Nature Reviews Microbiology. 2019;17(5):271–283.

2. Baross JA, Hoffman SE. Submarine hydrothermal vents and associated gradient environments as sites for the origin and evolution of life. Origins of Life and Evolution of the Biosphere. 1985;15(4):327–345.

3. Damer B, Deamer D. The hot spring hypothesis for an origin of life. Astrobiology. 2020;20(4):429–452.

4. Martin W, Russell MJ. On the origin of biochemistry at an alkaline hydrothermal vent. Philosophical Transactions of the Royal Society B: Biological Sciences. 2007;362(1486):1887–1926.

5. Leins A, Bregnard D, Vieth-Hillebrand A, Junier P, Regenspurg S. Dissolved organic compounds in geothermal fluids used for energy production: a review. Geothermal Energy. 2022;10(1):9.

6. Harold L. Drake KK, Carola Matthies. *Acetogenic prokaryotes*. vol 2. The prokaryotes. Springer Science; 2006:354–420.

7. Schuchmann K, Müller V. Autotrophy at the thermodynamic limit of life: a model for energy conservation in acetogenic bacteria. Nature Reviews Microbiology. 2014;12(12):809–821.

8. Widdel F, Pfennig N. A new anaerobic, sporing, acetate-oxidizing, sulfate-reducing bacterium, *Desulfotomaculum* (emend.) *acetoxidans*. Archives of Microbiology. 1977;112:119–122.

9. Spormann AM, Thauer RK. Anaerobic acetate oxidation to CO_2_ by *Desulfotomaculum acetoxidans*: Isotopic exchange between CO_2_ and the carbonyl group of acetyl-CoA and topology of enzymes involved. Archives of microbiology. 1989;152:189–195.

10. Londry K, Jahnke L, Des Marais D. Stable carbon isotope ratios of lipid biomarkers of sulfate-reducing bacteria. Applied and Environmental Microbiology. 2004;70(2):745–751.

11. Mardanov AV, Beletsky AV, Kadnikov VV, Slobodkin AI, Ravin NV. Genome analysis of *Thermosulfurimonas dismutans*, the first thermophilic sulfur-disproportionating bacterium of the Phylum *Thermodesulfobacteria*. Front Microbiol. 2016;7:950. doi:10.3389/fmicb.2016.00950

12. Moussard H, L’Haridon S, Tindall BJ, et al. *Thermodesulfatator indicus* gen. nov., sp. nov., a novel thermophilic chemolithoautotrophic sulfate-reducing bacterium isolated from the Central Indian Ridge. Int J Syst Evol Microbiol. Jan 2004;54(Pt 1):227–233. doi:10.1099/ijs.0.02669-0

13. Hügler M, Sievert SM. Beyond the Calvin cycle: autotrophic carbon fixation in the ocean. Annual review of marine science. 2011;3:261–289.

14. Berg IA, Kockelkorn D, Ramos-Vera WH, et al. Autotrophic carbon fixation in archaea. Nature Reviews Microbiology. 2010;8(6):447–460.

15. Mall A, Sobotta J, Huber C, et al. Reversibility of citrate synthase allows autotrophic growth of a thermophilic bacterium. Science. Feb 2 2018;359(6375):563-567. doi:10.1126/science.aao2410

16. Ivanovsky R, Sintsov N, Kondratieva E. ATP-linked citrate lyase activity in the green sulfur bacterium *Chlorobium limicola* forma thiosulfatophilum. Archives of Microbiology. 1980;128:239–241.

17. Möller D, Schauder R, Fuchs G, Thauer R. Acetate oxidation to CO_2_ via a citric acid cycle involving an ATP-citrate lyase: a mechanism for the synthesis of ATP via substrate level phosphorylation in *Desulfobacter postgatei* growing on acetate and sulfate. Archives of Microbiology. 1987;148:202–207.

18. Aoshima M, Ishii M, Igarashi Y. A novel enzyme, citryl□CoA synthetase, catalysing the first step of the citrate cleavage reaction in *Hydrogenobacter thermophilus* TK□6. Molecular microbiology. 2004;52(3):751–761.

19. Aoshima M, Ishii M, Igarashi Y. A novel enzyme, citryl□CoA lyase, catalysing the second step of the citrate cleavage reaction in *Hydrogenobacter thermophilus* TK□6. Molecular microbiology. 2004;52(3):763–770.

20. Hattori S, Galushko AS, Kamagata Y, Schink B. Operation of the CO dehydrogenase/acetyl coenzyme A pathway in both acetate oxidation and acetate formation by the syntrophically acetate-oxidizing bacterium *Thermacetogenium phaeum*. Journal of bacteriology. 2005;187(10):3471–3476.

21. Can M, Armstrong FA, Ragsdale SW. Structure, function, and mechanism of the nickel metalloenzymes, CO dehydrogenase, and acetyl-CoA synthase. Chemical reviews. 2014;114(8):4149–4174.

22. Nunoura T, Chikaraishi Y, Izaki R, et al. A primordial and reversible TCA cycle in a facultatively chemolithoautotrophic thermophile. Science. Feb 2 2018;359(6375):559-563. doi:10.1126/science.aao3407

23. Steffens L, Pettinato E, Steiner TM, et al. High CO_2_ levels drive the TCA cycle backwards towards autotrophy. Nature. Apr 2021;592(7856):784-788. doi:10.1038/s41586-021-03456-9

24. Nunoura T, Oida H, Miyazaki M, Suzuki Y. *Thermosulfidibacter takaii* gen. nov., sp. nov., a thermophilic, hydrogen-oxidizing, sulfur-reducing chemolithoautotroph isolated from a deep-sea hydrothermal field in the Southern Okinawa Trough. International journal of systematic and evolutionary microbiology. 2008;58(3):659–665.

25. Peretó JG, Velasco AM, Becerra A, Lazcano A. Comparative biochemistry of CO_2_ fixation and the evolution of autotrophy. Int Microbiol. Mar 1999;2(1):3–10.

26. Martin W, Russell MJ. On the origins of cells: a hypothesis for the evolutionary transitions from abiotic geochemistry to chemoautotrophic prokaryotes, and from prokaryotes to nucleated cells. Philos Trans R Soc Lond B Biol Sci. Jan 29 2003;358(1429):59-83; discussion 83-5. doi:10.1098/rstb.2002.1183

27. Peretó J. Out of fuzzy chemistry: from prebiotic chemistry to metabolic networks. Chem Soc Rev. Aug 21 2012;41(16):5394–403. doi:10.1039/c2cs35054h

28. Braakman R, Smith E. The emergence and early evolution of biological carbon-fixation. PLoS Comput Biol. 2012;8(4):e1002455.

29. Fuchs G. Alternative pathways of carbon dioxide fixation: insights into the early evolution of life? Annu Rev Microbiol. 2011;65:631–58. doi:10.1146/annurev-micro-090110-102801

30. Berg IA. Ecological aspects of the distribution of different autotrophic CO_2_ fixation pathways. Appl Environ Microbiol. Mar 2011;77(6):1925–36. doi:10.1128/aem.02473-10

31. Sánchez-Andrea I, Guedes IA, Hornung B, et al. The reductive glycine pathway allows autotrophic growth of *Desulfovibrio desulfuricans*. Nature communications. 2020;11(1):1–12.

32. Giovannelli D, Sievert SM, Hügler M, et al. Insight into the evolution of microbial metabolism from the deep-branching bacterium, *Thermovibrio ammonificans*. Elife. 2017;6:e18990.

33. Coleman GA, Davín AA, Mahendrarajah TA, et al. A rooted phylogeny resolves early bacterial evolution. Science. 2021;372(6542):eabe0511.

34. Vetriani C, Speck MD, Ellor SV, Lutz RA, Starovoytov V. *Thermovibrio ammonifican*s sp. nov., a thermophilic, chemolithotrophic, nitrate-ammonifying bacterium from deep-sea hydrothermal vents. International journal of systematic and evolutionary microbiology. 2004;54(1):175–181.

35. Sumi T, Harada K. Kinetics of the ancestral carbon metabolism pathways in deep-branching bacteria and archaea. Communications Chemistry. 2021;4(1):1–9.

36. Reysenbach A-L, Huber R, Stetter KO, et al. Phylum BI. Aquificae phy. nov. Bergey’s Manual® of Systematic Bacteriology. Springer; 2001:359–367.

37. Kawasumi T, Igarashi Y, Kodama T, Minoda Y. Isolation of strictly thermophilic and obligately autotrophic hydrogen bacteria. Agricultural and Biological Chemistry. 1980;44(8):1985–1986.

38. Kawasumi T, Igarashi Y, Kodama T, Minoda Y. *Hydrogenobacter thermophilus* gen. nov., sp. nov., an extremely thermophilic, aerobic, hydrogen-oxidizing bacterium. International Journal of Systematic and Evolutionary Microbiology. 1984;34(1):5–10.

39. Arai H, Kanbe H, Ishii M, Igarashi Y. Complete genome sequence of the thermophilic, obligately chemolithoautotrophic hydrogen-oxidizing bacterium *Hydrogenobacter thermophilus* TK-6. Journal of Bacteriology. 2010;192(10):2651–2652.

40. Fukuyama Y, Shimamura S, Sakai S, et al. Development of a rapid and highly accurate method for ^13^C tracer-based metabolomics and its application on a hydrogenotrophic methanogen. ISME communications. 2024;4(1):ycad006.

41. Shiba H, Kawasumi T, Igarashi Y, Kodama T, Minoda Y. The CO_2_ assimilation via the reductive tricarboxylic acid cycle in an obligately autotrophic, aerobic hydrogen-oxidizing bacterium, *Hydrogenobacter thermophilus*. Archives of microbiology. 1985;141(3):198–203.

42. Yamamoto M, Arai H, Ishii M, Igarashi Y. Characterization of two different 2-oxoglutarate: ferredoxin oxidoreductases from *Hydrogenobacter thermophilus* TK-6. Biochemical and biophysical research communications. 2003;312(4):1297–1302.

43. Ikeda T, Ochiai T, Morita S, et al. Anabolic five subunit-type pyruvate: ferredoxin oxidoreductase from *Hydrogenobacter thermophilus* TK-6. Biochemical and biophysical research communications. 2006;340(1):76–82.

44. Miura A, Kameya M, Arai H, Ishii M, Igarashi Y. A soluble NADH-dependent fumarate reductase in the reductive tricarboxylic acid cycle of *Hydrogenobacter thermophilu*s TK-6. Journal of bacteriology. 2008;190(21):7170–7177.

45. Aoshima M. Novel enzyme reactions related to the tricarboxylic acid cycle: phylogenetic/functional implications and biotechnological applications. Applied microbiology and biotechnology. 2007;75(2):249–255.

46. Li F, Hagemeier CH, Seedorf H, Gottschalk G, Thauer RK. *Re*-citrate synthase from *Clostridium kluyveri* is phylogenetically related to homocitrate synthase and isopropylmalate synthase rather than to *Si*-citrate synthase. J Bacteriol. Jun 2007;189(11):4299–304. doi:10.1128/jb.00198-07

47. Marco-Urrea E, Paul S, Khodaverdi V, et al. Identification and characterization of a *Re*-citrate synthase in *Dehalococcoides* strain CBDB1. Journal of bacteriology. 2011;193(19):5171–5178.

48. Lenz H, Buckel W, Wunderwald P, et al. Stereochemistry of si□Citrate Synthase and ATP□Citrate□Lyase Reactions. European journal of biochemistry. 1971;24(2):207–215.

49. Verschueren KH, Blanchet C, Felix J, et al. Structure of ATP citrate lyase and the origin of citrate synthase in the Krebs cycle. Nature. 2019;568(7753):571-575.

50. Shiba H, Kawasumi T, Igarashi Y, Kodama T, Minoda Y. Effect of organic compounds on the growth of an obligately autotrophic hydrogen-oxidizing bacterium, Hydrogenobacter thermophilus TK-6. Agricultural and biological chemistry. 1984;48(11):2809–2813.

51. Feng X, Tang K-H, Blankenship RE, Tang YJ. Metabolic flux analysis of the mixotrophic metabolisms in the green sulfur bacterium *Chlorobaculum tepidum*. Journal of Biological Chemistry. 2010;285(50):39544–39550.

52. Brandis-Heep A, Gebhardt NA, Thauer RK, Widdel F, Pfennig N. Anaerobic acetate oxidation to CO_2_ by *Desulfobacter postgatei*: I. Demonstration of all enzymes required for the operation of the citric acid cycle. Archives of microbiology. 1983;136:222–229.

53. Gebhardt NA, Linder D, Thauer RK. Anaerobic acetate oxidation to CO_2_ by *Desulfobacter postgatei*: 2. Evidence from ^14^C-labelling studies for the operation of the citric acid cycle. Archives of microbiology. 1983;136:230–233.

54. Makarova KS, Koonin EV. Filling a gap in the central metabolism of archaea: prediction of a novel aconitase by comparative-genomic analysis. FEMS microbiology letters. 2003;227(1):17–23.

55. Tsukahara K, Kita A, Nakashimada Y, Hoshino T, Murakami K. Genome-guided analysis of transformation efficiency and carbon dioxide assimilation by *Moorella thermoacetica* Y72. Gene. 2014;535(2):150–155.

56. Fuchs G, Stupperich E, Thauer RK. Acetate assimilation and the synthesis of alanine, aspartate and glutamate in *Methanobacterium thermoautotrophicum*. Archives of Microbiology. 1978;117:61–66.

57. d’Ippolito G, Dipasquale L, Fontana A. Recycling of Carbon Dioxide and Acetate as Lactic Acid by the Hydrogen□Producing Bacterium *Thermotoga neapolitana*. ChemSusChem. 2014;7(9):2678–2683.

58. Kim M, Le H, McInerney MJ, Buckel W. Identification and characterization of *Re*-citrate synthase in *Syntrophus aciditrophicus*. Journal of bacteriology. 2013;195(8):1689–1696.

59. Berg IA, Pettinato E, Böhnert P. Succinyl-CoA: acetate CoA-transferase functioning in the oxidative tricarboxylic acid cycle in Desulfurella acetivorans. Frontiers in Microbiology. 2022:4961.

60. Pierce E, Xie G, Barabote RD, et al. The complete genome sequence of *Moorella thermoacetica* (f. Clostridium thermoaceticum). Environmental microbiology. 2008;10(10):2550–2573.

61. Kim K, Chiba Y, Kobayashi A, Arai H, Ishii M. Phosphoserine phosphatase is required for serine and one-carbon unit synthesis in *Hydrogenobacter thermophilus*. Journal of bacteriology. 2017;199(21):e00409–17.

62. Hall TA. BioEdit: a user-friendly biological sequence alignment editor and analysis program for Windows 95/98/NT. Oxford; 1999:95–98.

63. Katoh K, Standley DM. MAFFT multiple sequence alignment software version 7: improvements in performance and usability. Molecular biology and evolution. 2013;30(4):772–780.

64. Nguyen L-T, Schmidt HA, Von Haeseler A, Minh BQ. IQ-TREE: a fast and effective stochastic algorithm for estimating maximum-likelihood phylogenies. Molecular biology and evolution. 2015;32(1):268–274.

