## Supporting information for "Impact of acetate on CO_2_ fixation pathways in thermophilic and hydrogenotrophic bacteria"

**Affiliations**

**Identification of *Re*-type citrate synthase (CS) in *Thermodesulfatator indicus*.**

CSs are classified into *Si*- and *Re*-types based on the stereospecificity of the binding manner of acetyl-coenzyme A (CoA) and oxaloacetate (OAA) (Fig. S2) and these two types do not share significant amino-acid sequence similarity [46]. ATP-citrate lyase (ACL) and citryl-CoA lyase (CCL) appear to be derived from *Si*-CS [45,48] and are therefore expected to exhibit same stereochemistry [23,45].

A relative of *Thermodesulfatator indicus*, *Thermosulfurimonas dismutans*, was reported to lack the gene for CS [11]; however, we found that *T. indicus* and *T. dismutans* harbors a putative *Re*-CS gene (Thein_0064 and ENJ40_01645), which has 52 and 53% identity to the *Re*-CS in *Clostridium kluyveri* [46], respectively while they lack *Si*-CS candidate genes. The phylogenic position of Thein_0064 was determined to more closely estimate its function because the amino acid sequence of *Re*-CS is quite similar to that of isopropylmalate synthase [46]. The analysis revealed that Thein_0064 formed a clade with biochemically confirmed *Re*-CS in *C. kluyveri* and *Syntrophus aciditrophicus* [58] and uninvestigated *Re*-CS homologues, with the grouping supported by a 100% bootstrap value (Fig. S3). Notably, this clade did not include proteins experimentally confirmed to have enzymatic activities other than CS. *T. indicus* also has a protein (Thein_0538) that was included in a clade with experimentally confirmed isopropylmalate synthases. *T. indicus* lacks candidate genes for ACL, CCL and citryl-CoA synthetase (CCS) in addition to *Si*-CS. Accordingly, we concluded that that *T. indicus* likely uses *Re*-CS for converting acetyl-CoA and OAA to citrate.

**Genes related to the reductive (r)TCA cycle and Wood Ljungdahl (WL) pathway in *T. indicus*, *T. ammonificans* and *H. thermophilus*.**

*T. ammonificans* and *H. thermophilus* encode a complete set of the genes for the TCA cycle (Table S2 and Fig. S1) [32,39]. In contrast, a BLASTp search suggested that the TCA cycle in *T. indicus* is incomplete due to the absence of genes for succinate dehydrogenase and succinyl-CoA synthetase (Table S2 and Fig. S1). Although some organisms use succinyl-CoA:acetate CoA transferase instead of succinate dehydrogenase [59], a CoA-transferase homolog was not found in the *T. indicus* genome.

BLASTp homology searches were also conducted against the estimated protein sequences of *T. indicus* using the WL pathway enzymes of *Moorella thermoacetica* [60] as queries. This analysis revealed that *T. indicus* possesses homologs of all WL pathway genes (E-values of less than $10^{-3}$, Table S3).

Taken together, the comparative genetic analyses suggested that the three organisms utilize the following metabolic pathways (Fig. S1). *T. indicus* has a complete WL pathway and an incomplete TCA cycle. The WL pathway likely fixes $CO_2$ into acetyl-CoA, which is further carboxylated to OAA through a part of the TCA cycle. Citrate and 2-oxoglutarate (2-OG) may be synthesized through the oxidative direction of the TCA cycle, and no connection between fumarate and succinyl-CoA was detected. In contrast to *T. indicus*, *Thermovibrio ammonificans* and *Hydrogenobacter thermophilus* were predicted to fix $CO_2$ through the rTCA cycle.

**Distribution of the genes for the seven known $CO_2$ fixation pathways other than the WL pathway and rTCA cycle.**

A BLASTp search was conducted against the estimated protein sequences of *T. indicus*, *T. ammonificans*, and *H. thermophilus* using key enzymes in the Calvin cycle, 3-hydroxypropionate bicycle, 3-hydroxypropionate-4-hydroxybutyrate cycle, and dicarboxylate-4-hydroxybutyrate cycle [14], as queries. The homology search revealed that none of the three organisms possessed a complete gene set for any of these $CO_2$ fixation pathways (Table S6). The absence of a glycine cleavage system, which has the potential to function as a $CO_2$-fixing reductive glycine pathway, has been reported for *H. thermophilus* and *T. ammonificans* [61]. *T. indicus* possesses only one protein of the glycine cleavage system (Thein_0130, which has ~40% identity to *Escherichia coli* GcvH). These results indicate that $CO_2$ is likely incorporated into the three bacteria exclusively through the WL pathway and/or rTCA cycle.

**Predicted working direction of the WL pathway and TCA cycle based on $^{13}C$ labeling patterns**

The WL pathway converts two molecules of $CO_2$ into CO (light green circles, Fig. 1) and formate (magenta circles, Fig. 1), which are then fixed to the C-1 and C-2 carbons, respectively, of the acetyl group of acetyl-CoA. Acetyl-CoA is expected to be carboxylated to pyruvate by pyruvate:ferredoxin oxidoreductase (POR), and pyruvate is further carboxylated to OAA by pyruvate carboxylase. Through this two-step carboxylation process, one $CO_2$ molecule is fixed to the C-1 carbon of pyruvate (and OAA; gray circles labeled "A" in Fig. 1) and the other is fixed to the C-4 carbon of OAA (gray circles labeled B in Fig. 1). Therefore, if $^{13}CO$ or $^{13}C$-formate is directly incorporated in the cells through the WL pathway, Ala synthesized from pyruvate should contain $^{13}C$ in either the C-2 or C-3 position, but not at C-1 (Fig. 1B). In other words, C-1 labeling of Ala in these experimental conditions indicates that the $^{13}C$-labeled CO and/or formate was first oxidized to $CO_2$ and then incorporated into the cells (Fig. S5, gray dot arrows).

In the rTCA cycle, acetyl-CoA is also carboxylated to OAA, as described above. OAA is converted into succinyl-CoA through four enzymatic reactions and the three intermediate substrates malate, fumarate, and succinate, and the generated succinyl-CoA is then carboxylated to 2-OG. The $CO_2$ fixed through carboxylation corresponds to the C-1 carbon of 2-OG and the product Glu (gray circles labeled "C" in Fig. 1). The two symmetrical intermediates of succinate and fumarate result in the shuffling of the C-1 and C-4, and C-2 and C-3 carbons of OAA during conversion to 2-OG, and therefore, two labeling patterns are predicted in Glu. 2-OG is further carboxylated into isocitrate, and the $CO_2$-derived carbon forms a carboxyl group bound to the C-3 carbon on the main chain (circles labeled "D" in Fig. 1 and Fig. S6). As described above, *Si*- and *Re*-CS cleave citrate into acetyl-CoA and OAA in different ways. In the case of *Si*-CS, two carbons fixed into succinyl-CoA (circles labeled "C" and "D") form the C-1 and C-4 carbons of

OAA (Fig. 1, Fig. S6A). In the case of *Re*-CS, the two carbons fixed into succinyl-CoA (circles labeled "C" and "D") form the C-1 carbons of OAA and acetyl-moiety of acetyl-CoA, with the C-1 carbon of acetyl-CoA corresponding to the C-2 carbon of pyruvate (Fig. 1, Fig. S6B).

The working direction of the TCA cycle under chemolithomixotrophic conditions can be determined based on labeling patterns obtained with $^{13}C_2$ acetate as a tracer. Here, we considered three possible TCA cycle variants based on gene sets for TCA cycle genes in *T. indicus*, *T. ammonificans*, and *H. thermophilus* (Fig. 1A). The first possible variant is a rTCA cycle in which the net flux moves in the reductive direction (anticlockwise direction in Fig. 1A). The second variant is the oxidative (o)TCA cycle, which works in the oxidative direction (clockwise direction in Fig. 1A). The third variant is an incomplete TCA cycle in which the connection between 2-OG and OAA is absent.

When the rTCA cycle is active, the incorporated $^{13}C_2$ acetate is converted into $^{13}C_2$ citrate via acetyl-CoA (Fig. S6). The $^{13}C_2$ citrate cleaved by *Si*-CS or its homologs (CS domain in ACL or CCL) forms $^{13}C_1$ acetyl-CoA and $^{13}C_1$ OAA (Fig. S6A). The detection of $^{13}C_1$ alanine (Ala) and $^{13}C_1$ aspartate (Asp) derived from $^{13}C_1$ acetyl-CoA and $^{13}C_1$ OAA satisfies the operation of the rTCA cycle with *Si*-type citrate cleavage. In contrast, $^{13}C_0$ acetyl-CoA and $^{13}C_2$ OAA are generated when $^{13}C_2$ citrate is cleaved by *Re*-CS (Fig. S6B). Therefore, the working direction of the TCA cycle cannot be determined based on labeling pattern of Ala and Asp in cells grown chemolithomixotrophically with acetate and an active *Re*-CS enzyme because the$^{13}C_2$ OAA (=$^{13}C_2$ Asp) obtained from citrate cannot be distinguished from that formed directly from $^{13}C_2$ acetate through acetyl-CoA.

When the oTCA cycle functions with active incorporation of $^{13}C_2$ acetate, $^{13}C_4$ Glu is detected as a result of $^{13}C_4$ citrate formation followed by $^{13}C_4$ 2-OG production (Fig. S7). In the case of an active oTCA cycle with *Si*-CS, $^{13}C_4$ Asp is produced via $^{13}C_4$ succinyl-CoA (Fig. S7A). In contrast, *Re*-CS forms $^{13}C_3$ succinyl-CoA and therefore, $^{13}C_3$ Asp will be detected (Fig. S7B). As the incomplete TCA cycle only provides $^{13}C_2$ Asp, as described below, detection of $^{13}C_4$ and

$^{13}C_3$ Asp indicates that the oTCA cycle is functioning with *Si*- and *Re*-CS-mediated reactions, respectively.

In the incomplete TCA cycle, 2-OG is formed by the decarboxylation of isocitrate, as in the case of the oTCA cycle (Fig. S8). Therefore, detection of $^{13}C_4$ glutamate (Glu) synthesized from $^{13}C_4$ 2-OG indicates the operation of either of the oTCA cycle or the incomplete TCA cycle, but not the rTCA cycle. Because the incomplete TCA cycle lacks the connection between 2-OG and OAA, OAA would be derived from pyruvate. Therefore, the detection of $^{13}C_4$ Glu and $^{13}C_2$ Asp and the absence of $^{13}C_4$ Asp and $^{13}C_3$ Asp indicate that the incomplete TCA cycle is active.

**Growth conditions**

*T. indicus* was cultivated statically in modified JCM354 medium at 70 °C under a gas mixture of 80% $H_2$ and 20% $CO_2$ (0.2 MPa). The modified JCM354 medium consisted of the inorganics of JCM354 medium and $NaHCO_3$. To prepare the modified medium, 3 mL of medium without $Na_2S$ and $NaHCO_3$ was autoclaved in a glass tube with a butyl rubber stopper under a 0.10 MPa $H_2$ atmosphere. $H_2$ was then added to the sealed tube at 0.16 MPa, followed by the addition of $CO_2$ at 0.20 MPa. Prior to inoculation, sterilized 5% (w/v) $Na_2S{\cdot}9H_2O$ solution (pH 8.0) and 10% (w/v) $NaHCO_3$ solution were added to the medium to give final concentrations of $Na_2S$ and $NaHCO_3$ of 0.05% (w/v) and 0.1% (w/v), respectively. For chemolithomixotrophic cultivation, sodium acetate (2 mM final concentration) was added to the inorganic-modified JCM 354 medium.

*T. ammonificans* was cultivated statically in an inorganic medium at 75 °C under a gas mixture of 80% $H_2$ and 20% $CO_2$ (0.2 MPa). Modified JCM 268 MJ(-N) synthetic seawater with lower concentrations of NaCl (2%) and $NH_4Cl$ (0.002%) was used as a basal medium. To prepare the medium, 4 mL of the basal medium was autoclaved in a glass tube with a butyl rubber atopper under a 0.10 MPa $H_2$ atmosphere. A gas mixture of 80% $H_2$ and 20% $CO_2$ was then added to the sealed tube at 0.20 MPa. Prior to inoculation, sterilized $Na_2S{\cdot}9H_2O$ and $KNO_3$

solution were added to the medium as described above. For chemolithomixotrophic cultivation, sodium acetate (10 mM final concentration) was added to the modified MJ(-N) synthetic seawater medium.

*H. thermophilus* TK-6 was cultivated in 5 mL of hydrogen-oxidizing bacteria (HOB) medium (see the Supporting Information for details) in a 100-mL glass serum bottle with a butyl rubber stopper at 70 °C under a gas phase (0.1 MPa) of 75% $H_2$, 10% $O_2$, and 15% $CO_2$ with shaking at 200 rpm. The HOB medium contained the following components per liter: 3 g $(NH_4)_2SO_4$, 1 g $KH_2PO_4$, 2 g $K_2HPO_4$, 0.5 g $K_2HPO_4$ g 0.25 g NaCl, 0.03 g $CaCl_2$, 0.014 g $FeSO_4 \cdot 7H_2O$, and 0.5 mL of a trace element solution reported previously [37]. For chemolithomixotrophic cultivation, sodium acetate (10 mM final concentration) was added to the inorganic medium.

**Cell cultivation for isotopologue and isotopomer analyses**

To examine $CO_2$ incorporation in *T. indicus* cells grown chemolithoautotrophically, cells were harvested 7 h after the addition of 10% (v/v of gas phase) $^{13}CO_2$ during the late exponential phase, as described below. After the gas phase ($H_2$:$CO_2$) of the late exponential culture was degassed to 0.1 MPa, 4 mL of $^{13}CO_2$ was injected into the culture tube and $H_2$ gas was added to 0.2 MPa. To examine the incorporation of $CO_2$ in *T. indicus* cells grown chemolithomixotrophically with acetate, $^{12}C$ sodium acetate (2 mM final concentration) and 2 mL of $^{13}CO_2$, which corresponded to 5% v/v of the final gas phase ($H_2$:$CO_2$) and 25% of all $CO_2$ in the test tube, was added prior to the cultivation. To examine the incorporation of acetate and formate in *T. indicus* cells, [1,2-$^{13}C_2$] sodium acetate or $^{13}C$ sodium formate (2 mM final concentration), respectively, was added to the inorganic medium prior to cultivation. To examine CO incorporation in *T. indicus* cells, 1 mL of $^{13}CO$ was added to the gas phase, which was pressurized by $H_2$ to 0.16 MPa, of the medium prepared chemolithoautotrophically, followed by the addition of $CO_2$ at 0.20 MPa to give a final CO:$CO_2$ ratio of 1:3.

To examine $CO_2$ incorporation in *T. ammonificans* cells grown chemolithoautotrophically, cells in the late exponentially phase were harvested 3 h after the addition of 10% $^{13}CO_2$ (v/v of the final gas phase), as described below. After the gas phase ($H_2$:$CO_2$) of the culture in the late exponential phase was degassed to 0.1 MPa, 2.4 mL of $^{13}CO_2$ was injected into the culture tube, and $H_2$ was then added to 0.2 MPa. To examine $^{13}CO$ incorporation in *T. ammonificans* cells, 2.4 mL of $^{13}CO$ was added to the $H_2$ atmosphere and a gas mixture of 80% $H_2$ and 20% $CO_2$ was added to 0.20 MPa. After a 16-h incubation, the cells were harvested by centrifugation. To examine acetate incorporation in *T. ammonificans* cells grown chemolithomixotrophically, [1,2-$^{13}C_2$] sodium acetate (10 mM final concentration) was added to the inorganic medium prior to cultivation. After a 16-h incubation, the cells were harvested by centrifugation.

To obtain labelled amino acids from *H. thermophilus* cells grown chemolithoautotrophically or chemolithomixotrophically, 10 mL of $^{13}CO_2$, which corresponded to ~10% (v/v) of the gas phase, was added to exponential phase cultures approximately 6 h after inoculation. Cells were harvested by centrifugation 2.5 h after the addition of $^{13}CO_2$. To examine the incorporation of acetate in *H. thermophilus* cells, [1,2-$^{13}C_2$] sodium acetate (10 mM final concentration) was added to the inorganic medium prior to cultivation, and cells were harvested by centrifugation in the late exponential phase.

**Proteomic analysis**

*T. indicus* and *H. thermophilus* cells grown chemolithoautotrophically and chemolithomixotrophically with acetate were used for shotgun proteomic analysis, as described previously [22] with minor modifications. Cells stored at −80 °C were suspended in 400 µl cell lysis buffer (100 mM triethylammonium bicarbonate (TEAB; pH 8.6) and 2 mM phenylmethylsulfonyl fluoride (PMSF)) and were then disrupted by sonication. The protein concentration of the cell-free extract was determined fluorometrically using a Qubit fluorometer (Thermo Fisher Scientific). Cell extracts corresponding to 10 µg protein were evaporated to dryness and were then dissolved in denaturing buffer consisting of MPEX PTS Reagent Solutions A and B (GL Science, Tokyo, Japan). The denatured proteins were reduced by the addition of dithiothreitol (25 mM final concentration; MS grade, Thermo Fisher Scientific) and incubation at 95 °C for 5 min, followed by room temperature incubation for 25 min. After the addition of iodoacetamide (25 mM final concentration; MS grade, Thermo Fisher Scientific), the resulting solutions were incubated at room temperature for 30 min for alkylation. Protein digestion was initiated by the addition of 100 ng trypsin protease (MS grade, Thermo Fisher Scientific), and the solution was incubated at 37 °C for 3 h before the proteins were further digested at 30 °C overnight by the subsequent addition of 100 ng trypsin protease. Detergents in the samples were then removed with MPEX PTS Reagent Solutions C and D (GL Science) according to the manufacturer's instructions. Residual surfactants were removed with a Pierce Detergent Removal Spin Column (Thermo Fisher Scientific). The digested protein solutions were evaporated to dryness, And the dried proteins were resuspended in 2% acetonitrile and 0.1% trifluoroacetic acid in water before being subjected to LC-MS/MS analysis, as described below.

Peptides were separated using an Ultimate 3000 RSLCnano nanoflow LC-system (Thermo Fisher Scientific) equipped with a reverse-phase column (Zaplous alpha Pep-C18, 3 μm, 120 A 0.1×150 mm; AMR Inc., Tokyo, Japan). A linear gradient of 5%-45% solvent B (100% acetonitrile) against solvent A (0.1% formic acid) was applied to the column over 100 min at a constant flow rate of 500 nl/min. via a nano-electrospray ion source (Dream Spray, AMR Inc.). The column temperature was set at 35 °C. An Orbitrap Fusion Tribrid mass spectrometer (Thermo Fisher Scientific) was operated in positive ion mode with an electrospray voltage of 1.5 kV and ion transfer tube temperature of 250 °C. Full scan MS spectra were acquired using an Orbitrap mass analyzer (*m/z* range: 350‑1800, resolution:120,000 FWHM) using internal calibration (EASY-IC). MS/MS spectra were acquired with an Ion Trap mass analyzer (m/z range: auto, scan rate: rapid) using collision-induced dissociation (CID) MS/MS fragmentation.

Proteins were identified using the Proteome Discoverer 2.2 software package (Thermo

Fisher Scientific). The acquired spectra were searched against the list of coding DNA sequences (CDSs) identified in the genomes of the tested strains using the SEQUEST search engine and the following search parameters: enzyme cleavage specificity for trypsin, allowing two missed cleavage sites, cysteine carbamidomethylation and methionine oxidation, and a maximum error tolerance for precursor and fragment masses were 10 ppm and 0.6 Da, respectively. Peptides corresponding to a <1% protein false discovery rate (FDR) were used in the calculations.
